# A modular toolbox for the optogenetic deactivation of transcription

**DOI:** 10.1101/2023.11.06.565805

**Authors:** Philipp Muench, Matteo Fiumara, Nicholas Southern, Davide Coda, Sabine Aschenbrenner, Bruno Correia, Johannes Gräff, Dominik Niopek, Jan Mathony

## Abstract

Light-controlled transcriptional activation is a commonly used optogenetic strategy that allows researchers to regulate gene expression with high spatiotemporal precision. The vast majority of existing tools are, however, limited to light-triggered induction of gene expression. Here, we inverted this mode of action and created two complementary optogenetic systems capable of efficiently terminating transcriptional activation in response to blue light. First, we designed highly compact regulators, by photo-controlling VP16 transactivation peptide exposure. Then, applying a two-hybrid strategy, we engineered LOOMINA (**l**ight **o**ff-**o**perated **m**odular **in**ductor of transcriptional **a**ctivation), a versatile transcriptional control platform for mammalian cells that is highly adaptable and compatible with various effector proteins. Leveraging the flexibility of CRISPR systems, we integrated LOOMINA with Cas9 as a DNA-binding domain to control transcription from various endogenous promoters with exceptionally high dynamic ranges in multiple cell lines, including neuron-like cells. Both functionally and mechanistically, LOOMINA represents a valuable addition to the optogenetic repertoire for transcriptional regulation.

## Introduction

Transcriptional control is a key strategy for regulating RNA and protein levels in cells. The ability to turn gene expression on or off in a user-defined manner provides tailored control over a wide variety of cellular processes. As a result, tools for the induction of gene expression have applications in all areas of biology from basic research to biotechnology. Among these approaches, optogenetic systems are particularly appealing due to the high degree of spatiotemporal control they offer.

Various optogenetic tools are available to regulate transcription in mammalian cells. The majority are based on pairs of protein domains that form heterodimers in response to light. Fusing one of the dimerization domains to a DNA-binding domain (DBD) and the other to a transactivation domain (TAD) enables light-induced recruitment of the TAD to DNA, as has been successfully demonstrated in cell culture^1–6^ and *in vivo*^7,8^.

The modularity of these two-hybrid systems allows for easy modifications by exchanging individual components. Existing tools include both blue-and red light-responsive dimerizers that have been combined with different DBDs and transactivator domains, depending on the application context^2,3,9–12^. In particular, the repurposing of CRISPR nucleases, such as Cas9, as DNA-binding modules has increased the versatility of these systems by enabling the simple and efficient targeting of endogenous loci^2,3,5^. For instance, recruitment of the VP64 or p65 transactivation domain to a catalytically dead Cas9 (dCas9) via fusions to the well-established CRY2/CIB1 photosensory heterodimerization system facilitates potent light-activated transcription^3,5^. In addition to TAD recruitment, several mechanistically distinct optogenetic CRISPR activation (CRISPRa) strategies have been described, including split-dCas9 architectures^2^, optogenetic recruitment of TADs to single guide RNAs (sgRNA)^2^, light-activated anti-CRISPR proteins^13,14^ or the steric control of Cas9 via light-dependent occlusion of the DNA-binding groove^15^. Importantly, all these strategies share the common feature of light-induced transcriptional activation.

However, if longer periods of active transcription are desired, an inverted mechanism of action, i.e. light-induced termination of transcriptional activation would be highly advantageous. This strategy could minimize the time that cells are exposed to light and allow for the controlled and rapid shutdown of gene expression upon illumination. Yet, only a few light-off systems for transcriptional regulation have been developed. While the activity of dCas9-VP64 could be successfully controlled via light-dependent subcellular re-localization, this indirect approach yielded only a modest fourfold difference in gene expression between the light and dark condition^4^. Alternative strategies showed higher dynamic ranges of gene expression control, but relied on sequence-specific DBDs, hence rendering the optogenetic control of selected, endogenous genes infeasible^16–18^.

To fill this gap in the optogenetic repertoire, we set out to create a versatile light-off toolkit for transcriptional control. First, building upon our previous work^19,20^, we photo-controlled the VP16 transcriptional transactivator peptide by incorporating it into the C-terminal Jɑ helix of the blue light-inducible LOV2 domain of *Avena sativa* phototropin 1 (*As*LOV2). Following this initial compact, single-component design, we created a powerful modular system based on a two-hybrid-like approach employing the LOVTRAP optogenetic dimerizer^21^. LOVTRAP consists of the *As*LOV2 domain and a peptide, Zdk (or Z-dark), which binds *As*LOV2 only in the dark. Fusing one or more *As*LOV2 domains to a DBD and linking Zdk to a transactivation domain facilitated robust activation of transcription in the dark and efficient deactivation in the light. Our **l**ight-**o**ff **o**perated **m**odular **in**ducer of transcriptional **a**ctivation (**LOOMINA**) is compatible with various DBDs including Gal4 and Cas9, and multiple TADs such as VP64, p65 or VPR, enabling the optogenetic expression control of reporter genes, as well as the induction from endogenous promoters. With up to 1000-fold changes in gene expression at endogenous loci, the system has an exceptionally wide dynamic range. Finally, the characterization of our toolbox with respect to spatial expression control as well as in different cell lines establishes LOOMINA as a robust and versatile means for the plug-and-play control of gene expression.

## Results

### Light-inhibited gene expression with photocaged transactivation peptides

To create a transcriptional control system that can be turned off by light, we first adapted a photocaging strategy that we had previously employed for optogenetic control of nuclear import and export of proteins^19,20^. The approach exploits the unique structural dynamics of the *As*LOV2 domain. In its dark-adapted state, the C-terminus of LOV2 constitutes a C-terminal α-helix, denoted Jα^22^, whereas photoinduction triggers the reversible unfolding of Jα, as well as its undocking from the LOV2 protein core^23,24^. As a transactivation domain, we used the widely applied *Herpes simplex* VP16 peptide, which adopts an α-helical structure when interacting with target proteins^25–27^. We reasoned that fusing VP16 to the Jɑ helix of *As*LOV2 would allow us to activate transcription in the dark, when Jɑ exhibits its helical fold, and deactivate VP16 in the relaxed light-adapted conformation of *As*LOV2. Of note, this light-off peptide control modality is essentially the reverse concept of previous, *As*LOV2-based peptide photocaging strategies employed by us and others^20,28,29^. As our initial design, we replaced the seven C-terminal amino acids of the *As*LOV2 domain with the 11 amino acid VP16 transactivation motif (**Fig. 1a**). Both sequences consist of negatively charged amino acids, interspersed with hydrophobic residues, rendering a successful incorporation into Jɑ highly likely.

**Fig. 1:**
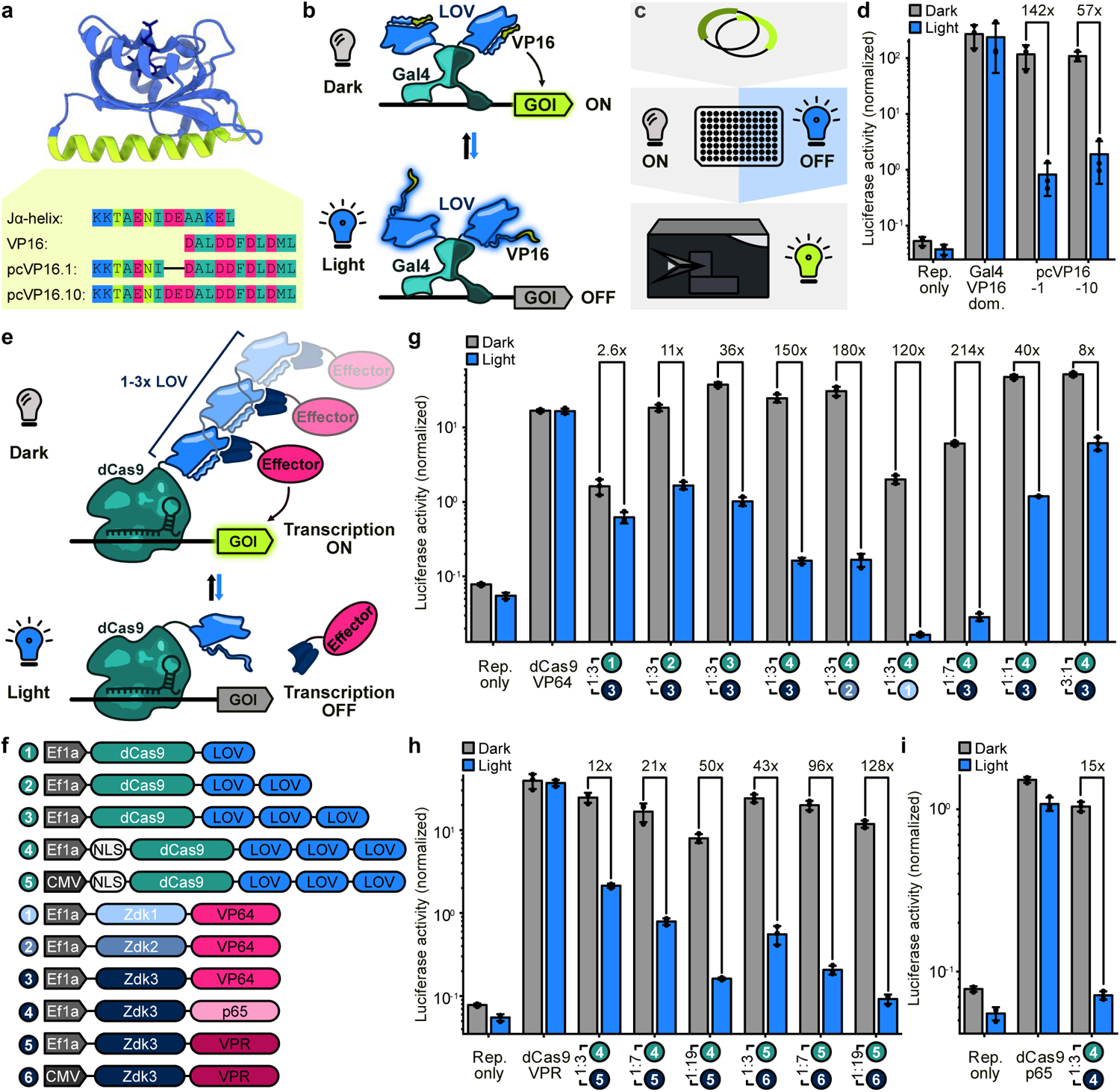
Development of potent light-inhibited transcriptional regulators. a Amino acid sequences of the pcVP16 lead candidates. The sequences corresponding to the C-terminus of the *As*LOV2 Jɑ helix and the VP16 transactivation peptide are depicted for comparison. **b** Schematic of transcriptional control with a photocaged Gal4-VP16. GOI: gene of interest. **c** Flowchart of the luciferase screening assay. **d** HEK293T cells were transfected with plasmids encoding (i) a *Renilla* luciferase, (ii) a Gal4-inducible firefly luciferase and (iii) the indicated Gal4-TAD fusions, followed by incubation of samples under blue light exposure or in the dark. The luciferase activity was measured 48 hours after transfection. Gal4-VP16-dom.: Gal4 fused to the complete VP16 transactivator domain. **e** Mechanism of action of the LOOMINA system. GOI: gene of interest. **f** Schematic of the constructs tested. **g-i** HEK293T cells were transfected with plasmids encoding (i) a firefly luciferase downstream of 13x TetO sites, (ii) a *Renilla* luciferase, (iii) dCas9 and (iv) VP64 (**g**), p65 (**h**) or VPR (**i**) as indicated by the circled numbers. As positive control a plasmid expressing a genetic fusion between dCas9 and the transactivator was used instead of the split constructs. The samples were either illuminated or kept in darkness, followed by measurements of luciferase activity after 48 h. **d**, **g-i** Bars represent the mean of three biological replicates. Data points indicate the individual biological replicates, which are the mean of three technical replicates. Error bars indicate S.D. Fold changes between illuminated and dark samples are indicated. Vector mass ratios between the dCas9-expressing construct and the transactivator construct are indicated for each sample. Experiments for **g-i** were performed together on a single plate and the same reporter only control was used as reference. Rep. only: reporter-only control.

This photo-controlled VP16 (pcVP16.1) variant was then fused to the C-terminus of a Gal4-DBD, to enable recruitment of the fusion protein to a Gal4-UAS reporter construct driving green fluorescent protein (GFP) expression (**Fig. 1b**). As benchmarks for gene induction potency and assay controls, we used a fusion of Gal4 to either the full-length VP16 domain or the VP16 transactivation peptide only. We then transiently co-transfected HEK293T cells with the GFP reporter and either the pcVP16.1 or the benchmark constructs and incubated the samples under blue light illumination (∼460 nm) or in the dark for 20 hours. Subsequent fluorescence measurements in a plate reader showed potent GFP expression for the control construct with the full-length VP16 domain, whereas the 11 amino acid-long VP16 peptide fused to Gal4 did not induce gene expression above basal levels (**Supplementary Fig. 1**). Our pcVP16.1 construct, in contrast, was both potent and light-switchable, exhibiting high reporter induction in the dark and only background levels of GFP expression when samples were incubated in the light. For further characterization, we switched to a more sensitive dual-luciferase assay and used a Gal4-UAS-driven firefly luciferase together with a co-transfected *Renilla* luciferase as internal reference (**Fig. 1c**). We compared pcVP16.1 with a number of alternative designs, again including Gal4-VP16-domain and Gal4-VP64 as benchmarks. pcVP16.1 exhibited a more than 140-fold induction of reporter gene expression in the dark as compared to the light condition (**Fig. 1d**, **Supplementary Fig. 2**). We also tested a naive fusion of the VP16 peptide to the untruncated Jα helix, which showed only modest light-dependency, when compared to pcVP16.1 (**Supplementary Fig. 2**). Extending the length of the *As*LOV2 controlled VP16 construct to include additional VP16 repeats, i.e. VP32-VP64, resulted in a graded increase in light-dependent activation, but similarly increased leaky expression in the light (**Supplementary Fig. 2b**).

Next, we fused pcVP16.1 to dCas9 (from *Streptococcus pyogenes*) as an alternative DBD that would allow the targeting of endogenous genomic loci. To this end, we used a firefly luciferase reporter, downstream of multiple Tet operator repeats and targeted a sgRNA to these operator sites in multiplex. The DBD exchange was generally successful, but longer VP48 or VP64 transactivator repeats were required to achieve sufficient induction, which came at the cost of increased levels of leaky expression in the light condition (**Supplementary Fig. 3**).

Thus, our reversed photocaging approach yielded a compact and potent transactivators with wide dynamic ranges of optogenetic gene expression control, but was only partially compatible with other DBDs.

### LOOMINA – a versatile transcription control platform based on the optogenetic LOVTRAP system

To create a light-off system for transcriptional regulation that is compatible with dCas9 and larger TADs such as VP64 and VPR, we took a different approach harnessing LOVTRAP, an *As*LOV2-based heterodimerization system thus far mainly employed to control protein subcellular relocalization^21,30^. In order to adapt LOVTRAP for direct transcriptional regulation, we fused the *As*LOV2 domain C-terminally to dCas9. Zdk2, a *As*LOV-binding peptide that dissociates upon illumination, was linked to the C-terminus of VP64 and expressed from a different plasmid^21^. We tested our strategy with the previously described luciferase reporter in HEK293T cells. Indeed, we observed light-dependent transcriptional activation with an up to 11-fold change in reporter activity between the light and dark samples (**Supplementary Fig. 4**). This effect was most pronounced when the transcriptional activator was added in excess of the dCas9-LOV encoding construct. Although this initial attempt was successful overall, the transcriptional activation in the dark was in all cases significantly lower (>26-fold) in comparison to the positive control samples expressing a direct fusion of dCas9 to VP64 (dCas9-VP64). Hypothesizing that domain architecture and order might affect the dimerization and transactivation efficiency, we screened N-terminal Zdk2-VP64 fusions and previously described alternative Zdk peptides (Zdk1 and Zdk3) under the same conditions as our prototype construct. Rearranging the domain order resulted in significantly improved transcriptional activation, especially for Zdk1, which was indistinguishable from the dCas9-VP64 benchmark in the dark (**Supplementary Fig. 5**). However, only small differences were observed between light and dark conditions, i.e. these designs showed impaired switchability. While the dynamic range of gene expression control could be improved by simply increasing the amount of dCas9-*As*LOV2 relative to the Zdk1-VP64 construct supplied during transfection, this improvement came at the cost of overall lower reporter activation in the dark condition (**Supplementary Fig. 6**). Therefore, we decided to take an alternative route of optimization. We hypothesized that an increase in the number of (light-dependent) Zdk-binding sites per Cas9 molecule could improve affinity and hence transactivator recruitment in the dark without compromising photo-switchability. To realize this concept, we concatenated up to three *As*LOV2 domains via genetic fusion and appended them to dCas9 via flexible glycine-serine linkers (**Fig. 1e-f**). Screening these constructs using our luciferase assay, we observed greatly increased differences between dark and light conditions from 2.6-fold for the single *As*LOV2 design to 36-fold when three *As*LOV2 domains were concatenated (**Fig. 1g**). We were able to further improve the system’s performance by adding multiple nuclear localization signals (NLS) to dCas9-3x*As*LOV2 (consisting of two SV40 NLS and a C-myc NLS), instead of the single SV40 NLS that was used up until this point. This new NLS configuration greatly reduced leaky reporter expression under illumination while maintaining full induction in the dark, resulting in a 150-fold change in reporter activity (**Fig. 1f-g**). We refer to this system and its derivatives as LOOMINA (for **l**ight-**o**ff **o**perated **m**odular **in**ducer of transcriptional **a**ctivation).

To investigate the modularity of LOOMINA, we explored the effects of altering its LOVTRAP components. We combined VP64 separately with different Zdk variants (Zdk1-3), that were reported to have varying affinities to *As*LOV2^21^. Comparable differences between light and dark conditions were observed for all three Zdk variants tested (**Fig. 1g**, **Supplementary Fig. 7**). The combination of LOOMINA with Zdk1 resulted in exceptionally low background reporter activity in the light, but also in lower levels of induction in the dark (**Fig. 1g**, **Supplementary Fig. 7**). This observation reflects the comparatively lower affinity of Zdk1 to *As*LOV2 compared to Zdk2/3, as previously reported^21^. Zdk3 performed robustly under all tested conditions and was used in the further experiments. Similar to the Zdks, also the *As*LOV2 domain could be replaced by a recently published circularly permuted variant (cpLOV2)^31^, which performed comparably well (**Supplementary Fig. 8**).

Having identified a robust LOOMINA design, we next combined it with TADs other than VP64. As expected, Zdk3-VP64 could be readily replaced by Zdk3-VPR or Zdk3-p65 resulting in light-switchable transcriptional activation that was in the range of the dCas9-TAD positive controls in the dark and at basal levels in the light (**Fig. 1h-i**). In addition, the efficiency of LOOMINA could be tuned by varying the vector mass ratio of the transfected DBD and TAD components (**Fig. 1h-i**). In the case of VPR, for instance, lower DBD:TAD vector mass ratios (1:7 instead of 1:3) turned out beneficial (**Fig. 1h**). We could even further enhance the system’s performance in conjunction with VPR by expressing its components from a CMV-promoter instead of the slightly weaker Ef1ɑ promoter^32^ (**Fig. 1h**).

Next, we tested the compatibility LOOMINA with different DBDs. To this end, we replaced dCas9 with Gal4 or TetR and evaluated these variants using the reporter constructs described above (**Supplementary Fig. 9a-b**). Various configurations including one or two *As*LOV2 domains and an N-terminal cpLOV fusion in combination with either Zdk3-VP64 or Zdk3-VPR were tested (**Supplementary Fig. 9c-d**). As expected, all variants were potently light-switchable, with the Gal4 constructs exhibiting a stronger light response than the TetR configuration.

Finally, to further simplify the use of LOOMINA, we combined its two components into a single vector. To this end, we designed a LOOMINA transgene co-expressing the Zdk3-TAD (encoded 5’) and 3xNLS-dCas9-3xLOV parts (encoded 3’) from the same mRNA using a P2A peptide strategy^33^. This particular configuration was chosen to prevent the unintended formation of constitutively active fusion proteins in case of insufficient “self-splicing” by the P2A sequence, which would emerge if the Zdk3-TAD part was located 3’. All P2A constructs successfully mediated strong light-dependent transcriptional activation when expressed under an Ef1ɑ promoter (**Supplementary Fig. 10**) and will enable users to easily combine a single plasmid version of LOOMINA with their sgRNA of choice.

### LOOMINA works efficiently in various cell types and enables spatially controlled gene expression activation

A key advantage of optogenetic tools is the precise spatial control of activation that can be achieved by illuminating only specific areas of a sample. To demonstrate this feature of LOOMINA, we employed a TetO-controlled mCherry reporter targeted by a TetO-specific sgRNA and used VPR as TAD. Following transient transfection of HEK293T cells, we covered half of the culture with a black photomask followed by illumination with blue light. Fluorescence microscopy revealed a clear difference in reporter expression levels between the illuminated and non-illuminated parts of the dish, showing that LOOMINA facilitates spatially confined gene expression (**Fig. 2a**).

**Fig. 2:**
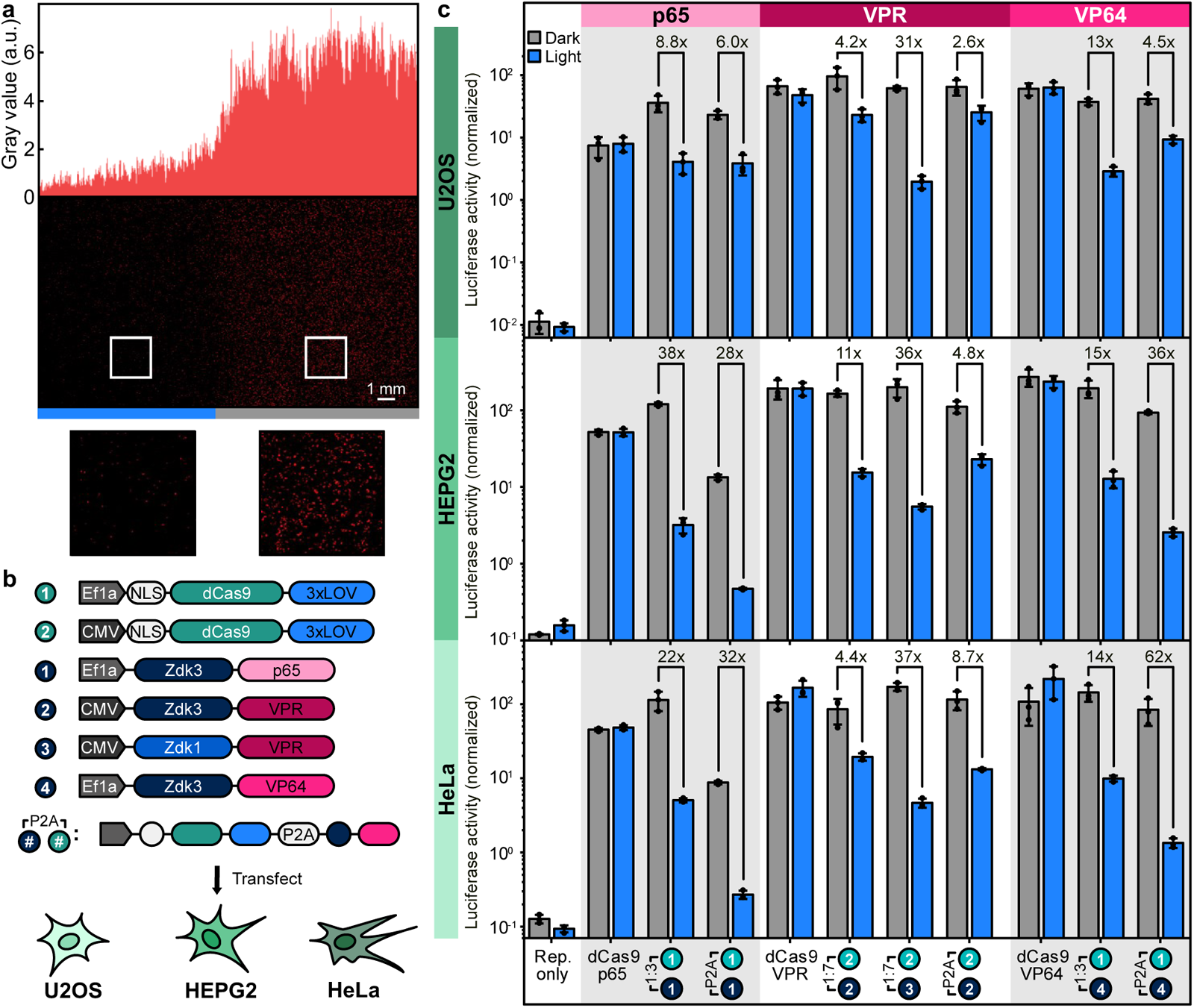
LOOMINA facilitates spatially confined gene expression and mediates robust transcriptional control in various cell lines. a. HEK293T cells were transfected with plasmids encoding (i) a tet-inducible mCherry reporter, (ii) a TetO-targeting sgRNA and (iii/iv) LOOMINA in the NLS-dCAs9-3xLOV/Zdk3-VPR configuration. Cells were illuminated through a photomask covering half of the plate for 48 h. Fluorescence microscopy images are shown along with a quantification of the fluorescent signal across the x-axis of the image. **b** Overview of the constructs and cell lines used. **c.** U2OS, HEPG2 or HeLa cells were transfected with plasmids encoding (i) a firefly luciferase downstream of 13x TetO sites, (ii) a *Renilla* luciferase, (iii) the LOOMINA constructs as indicated by the circled numbers. As positive control a plasmid expressing a genetic fusion between dCas9 and the transactivator was used instead of the split constructs. Samples were either illuminated or kept in the dark, followed by measurements of luciferase activity after 48 h. Bars represent means of three biological replicates. Data points indicate individual biological replicates, which are the mean of three technical replicates. Error bars indicate S.D. Fold changes between illuminated and dark samples are indicated. Rep. only: reporter-only control.

To further assess LOOMINA’s applicability, we tested the system in several commonly used human cell lines such as HeLa, U2OS, HepG2. Evaluation of our lead candidate 3xNLS-dCas9-3xLOV in combination with all three TADs (VP64, p65 and VPR) via luciferase assays revealed light-dependent transcriptional activation in each cell line under all tested conditions (**Fig. 2b and c**). For every cell line, at least one TAD was identified that mediated a more than 30-fold switch in reporter activity between dark and light conditions (**Fig. 2c**). In addition, all constructs achieved activation levels in the dark that were comparable to the respective dCas9-TAD fusion protein benchmarks. Importantly, assay conditions were not individually optimized for each cell line or TAD, i.e. robust photoswitching was readily obtained simply by using LOOMINA with the same configuration and vector mass ratios as previously in HEK293T cells.

### LOOMINA mediates strong transcriptional control at endogenous loci

One major limitation of previously published light-off switches for gene expression control is their reliance on sequence-specific DBDs^16–18^. As LOOMINA is compatible with multiple DBDs, fusion with dCas9 allows for site-specific, light-dependent transcriptional activation at targeted genomic loci. We investigated this feature by co-transfecting HEK293T or HeLa cells with plasmids encoding LOOMINA in combination with a mixture of sgRNA-expressing constructs that target the promoter region of a selected gene and measuring its expression by real-time quantitative PCR (RT-qPCR) three days post transfection (**Fig. 3a**). We tested the LOOMINA VP64 and VPR variants, as well as the all-in-one P2A-constructs (**Fig. 3b**). Targeting the promoter of *IL1RN* in HEK293T cells resulted in blue light-inhibited control with exceptionally high dynamic ranges spanning two and three orders of magnitude for VP64 and VPR, respectively (**Fig. 3c-d**). To our knowledge this is the most effective *As*LOV2-based photoswitch reported in mammalian cells to date. When we targeted *IL1RN* in HeLa cells, the overall transcriptional activation reached by the dCas9-TAD fusion controls was lower, but LOOMINA again spanned the entire range from full activation in the dark to background levels of the negative control under light exposure (**Fig. 3e**). Similar results were obtained for the targeted activation of *MYOD* and *OCT4* in HEK293T cells (**Fig. 3f-g**), highlighting that LOOMINA enables tight transcriptional regulation of endogenous loci in a plug-and-play manner.

**Fig. 3:**
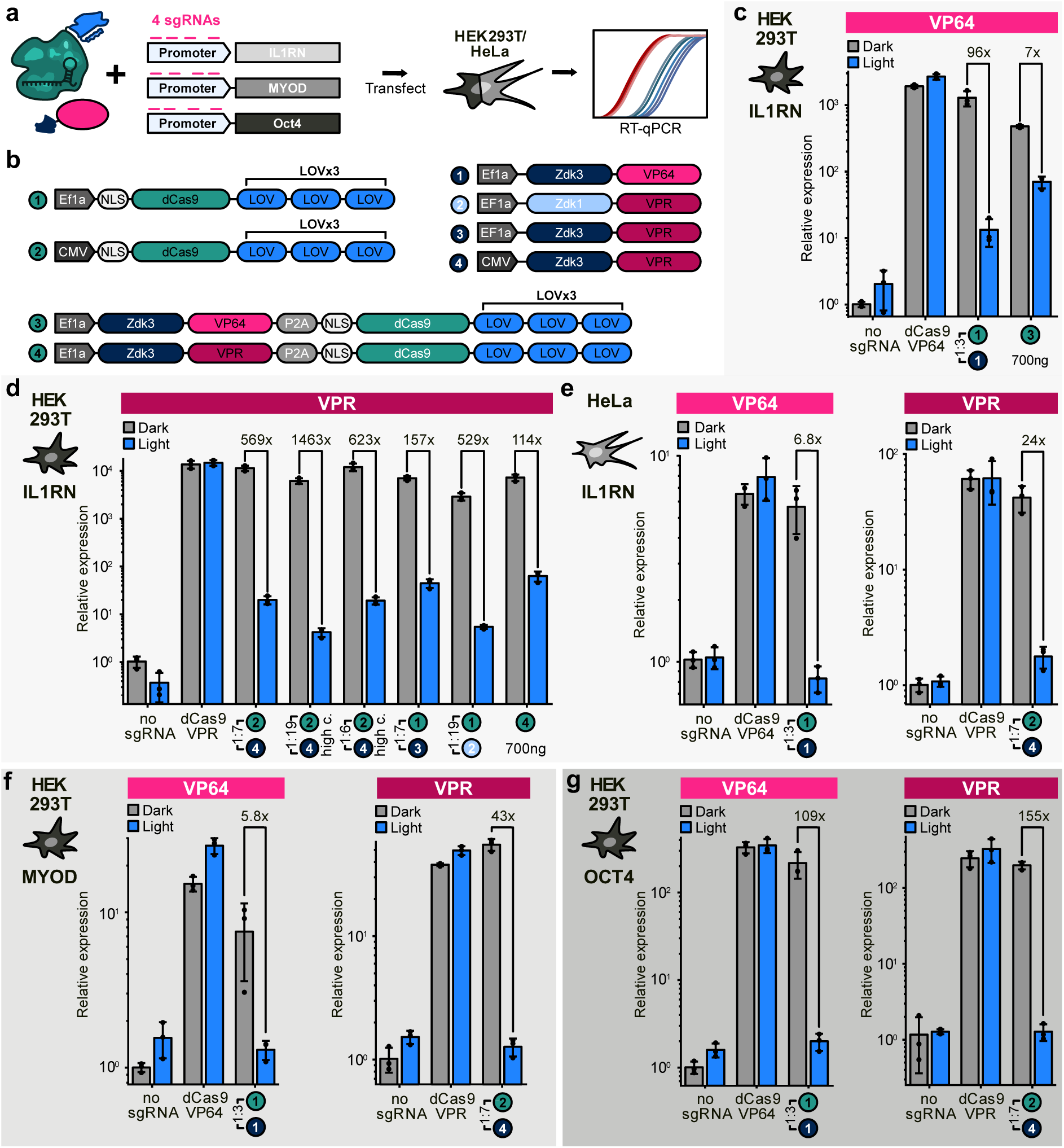
LOOMINA enables endogenous transcriptional control with high dynamic range. a. Overview of the experimental workflow for activating transcription at endogenous loci. **b** Schematics of the vectors that were used for the control of genomic promoters. **c-g** HEK293T or HeLa cells were co-transfected with (i) a cocktail of plasmids targeting different sites either in the promoter regions of *IL1RN* (**c-e**), *MYOD* (**f**) or *OCT4* (**g**) and (ii) plasmids encoding for the individual LOOMINA components. As a positive control, a plasmid expressing a fusion between dCas9 and either VP64 or VPR was used. The circled numbers refer to the combination of Cas9 and TAD used and correspond to the vectors outlined in panel **b**. The vector mass ratios between the dCas9-expressing construct and the transactivator construct are indicated. Samples were either illuminated or kept in the dark, followed by RT-qPCR analysis of gene expression after 72 hours. Bars represent the mean of three biological replicates. Data points indicate individual biological replicates, which are the mean of three technical replicates. Error bars indicate S.D. Fold changes between illuminated and dark samples are indicated. high c., 700 ng of the combined LOOMINA components were transfected instead of 400 ng.

To demonstrate LOOMINA’s robustness with respect to laboratory-specific experimental conditions and setups, we aimed to evaluate our system in a neuron-like cell line using a different illumination setup in another laboratory. In contrast to the experiments described above, we adopted a commercially available LED array and tested different dCas9-*As*LOV2 variants together with Zdk3-VP64 and Zdk3-VPR in Neuro2a (N2a) cells, a neuron-like cell line, using a luciferase reporter. Despite weaker overall reporter activation in the neuron-like cells as compared to the other cell lines tested, light-dependent gene expression control was obtained for the LOOMINA samples (**Supplementary Fig. 11**).

Next, moving from reporter control to optogenetic control of endogenous gene expression in N2a cells, we decided to use a shorter light pulsation cycle (1 s light on, 1 s light off), which reduced the leakiness of LOOMINA in the light (**Supplementary Fig. 12**). These experiments revealed an up to 5-fold difference in transcriptional activation when targeting the murine *Arc* locus, an important regulator of synaptic plasticity and memory^34^ (**Supplementary Fig. 12** and 13a**-b**). In order to cross-validate the RT-qPCR results, we performed an RNAscope^35^ experiment, using specific probes against the *Arc* mRNA. To visualize all components of the system, we added a fluorescent mCherry reporter to the Zdk3-VPR construct via a P2A peptide^33^ and co-expressed GFP from the sgRNA-encoding plasmid. We confirmed by RT-qPCR that the system’s functionality was not affected by the presence of mCherry or GFP (**Supplementary Fig. 13b**). Confocal image analysis revealed that 10–20% of cells co-expressed all transfected components, i.e. mCherry/Zdk3-VPR, dCas9-LOV and the sgRNAs (**Fig. 4a-b**, **Supplementary Fig. 13c-d**). Importantly, quantification of *Arc* mRNA levels in the triple-positive cells confirmed that *Arc* expression was significantly lower when cells were illuminated, confirming that LOOMINA induces transcription only in the dark (**Fig. 4c**).

**Fig. 4:**
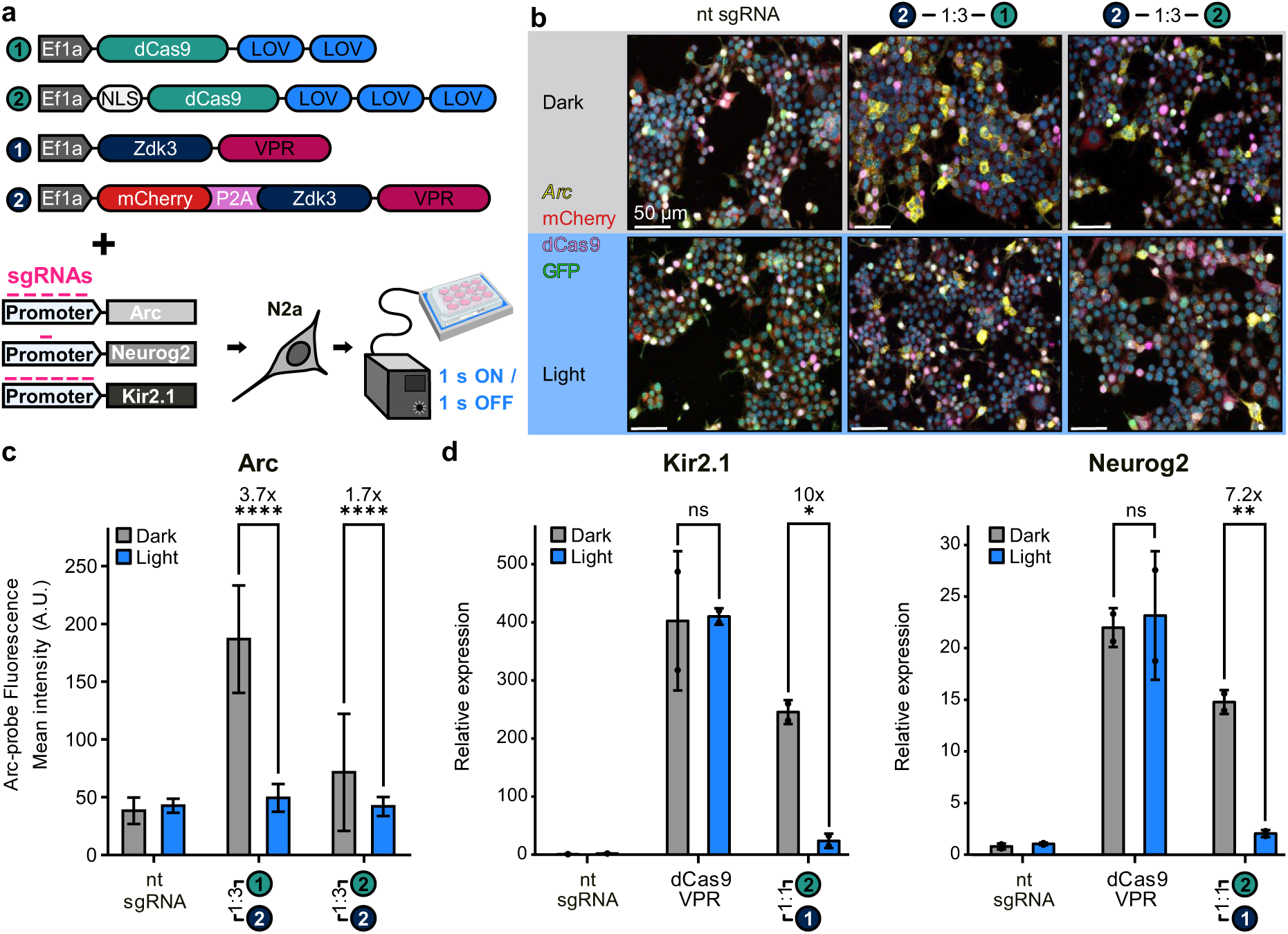
LOOMINA enables light-controlled expression of endogenous genes in a neuronal-like cell line. a. Schematic of the LOOMINA configurations used in N2a cells. Overview of the experimental workflow for the activation of endogenous transcription in N2a cells. The number of multiplexed sgRNAs for each target gene is indicated. **b** Cells were co-transfected with plasmids encoding (i) the respective sgRNAs and (ii) the indicated LOOMINA constructs, followed by incubation under light exposure or in darkness for 48 hours. Expression of dCas9 (pink), mCherry (red) and GFP (green) was assessed by combined immunocytochemistry. Arc was visualized using sequence-specific RNAscope probes. Representative confocal RNAscope images are shown. **c** Quantification of the mean fluorescence intensity of the Arc-probe from **b**. Bars represent the mean of individual values from single cells. The number of cells analyzed per condition is shown in **Supplementary** Fig. 13. **d** N2a cells were transfected with the lead LOOMINA constructs alongside sgRNA expressing plasmids targeting either the *Kir2.1* (left) or *Neurog2* (right). A direct dCas9-VPR fusion was used as a positive control. Samples were either illuminated or kept in darkness for 48 hours, followed by RT-qPCR analysis of gene expression. Bars represent the mean of two biological replicates. Data points represent the individual biological replicates, which are the mean of three technical replicates. **b-d** Vector mass ratios between the dCas9-expressing construct and the transactivator construct are indicated. Circled numbers refer to the vectors outlined in panel **a**. Error bars indicate S.D. Fold changes between illuminated and dark samples are indicated. Statistical significance was assessed by two-way ANOVA with Sidak correction for multiple comparisons; n.s.: not significant; *p-value < 0.05; **p-value < 0.01, ****p-value<0.0001. nt sgRNA: non-targeting sgRNA control.

Finally, we evaluated the LOOMINA lead candidate design (NLS-dCas9-3xLOV, Zdk3-VPR) on additional endogenous loci in N2a cells, targeting the promoter regions of *Kir2.1* and *Neurog2*, genes implicated in synaptic transmission and neuronal differentiation, respectively^36,37^. In line with our previous results, LOOMINA significantly reduced *Kir2.1* and *Neurog2* transcriptional activation in the light while maintaining high expression levels comparable to the constitutive dCas9-VPR control in the dark state (**Fig. 4d**). Interestingly, in the case of *Neurog2*, high levels of activation were reached using only a single promoter-targeting sgRNA in contrast to multiplexed sgRNAs used in previous experiments and that are common for this type of assay^2,3,5,38^. Taken together, these findings underscore the ability of the LOOMINA system to provide custom regulation of endogenous gene expression, in this case in a neuron-like cell line.

## Discussion

Our work presents novel approaches for the light-off operated control of transcription represented by two complementary strategies, the miniaturized pcVP16 and the multi-purpose LOOMINA system (**Fig. 5**). The reverse-photocaging approach was inspired by concepts developed by us and others for the optogenetic control of diverse cellular processes, such as nuclear protein import^19,39^ or export^20,40^, protein degradation^41,42^ and protein-protein interactions^43^. In line with previous work, we modified the Jɑ helix of *As*LOV2 only from I539 onwards to preserve its function. Within the engineered portion, various different configurations were tolerated and functional, confirming that the C-terminal part of the Jɑ-helix is highly amenable to modifications (**Supplementary** Fig. 2). As a key novelty of our pcVP16 design, we have successfully fused the Jɑ helix to a structured helical motif, thereby inverting the previously described mode of LOV2-based peptide photocaging. While other approaches have relied on masking of signal peptides in the dark state, VP16 is active in the helical conformation and was in turn deactivated by Jɑ unfolding in the light. We expect that this strategy could easily be adapted for the optogenetic control of other helical peptides. Another remarkable difference between pcVP16 and previous systems is its highly potent light response with >100-fold changes in reporter expression (**Fig. 1d**), whereas other photocaging tools have exhibited only up to 40-fold changes in activity. Further investigation is needed to understand whether this particularly potent switchability of pcVP16 is due to specific features of the LOV2-VP16 fusion or a general property of the reverse photocaging concept developed here. Finally, we note that pcVP16 is particularly compact. Including Gal4 as DBD, the complete fusion protein is only 318 amino acids long, rendering it an interesting candidate for future control of virally delivered transgenes *in vivo*.

**Fig. 5:**
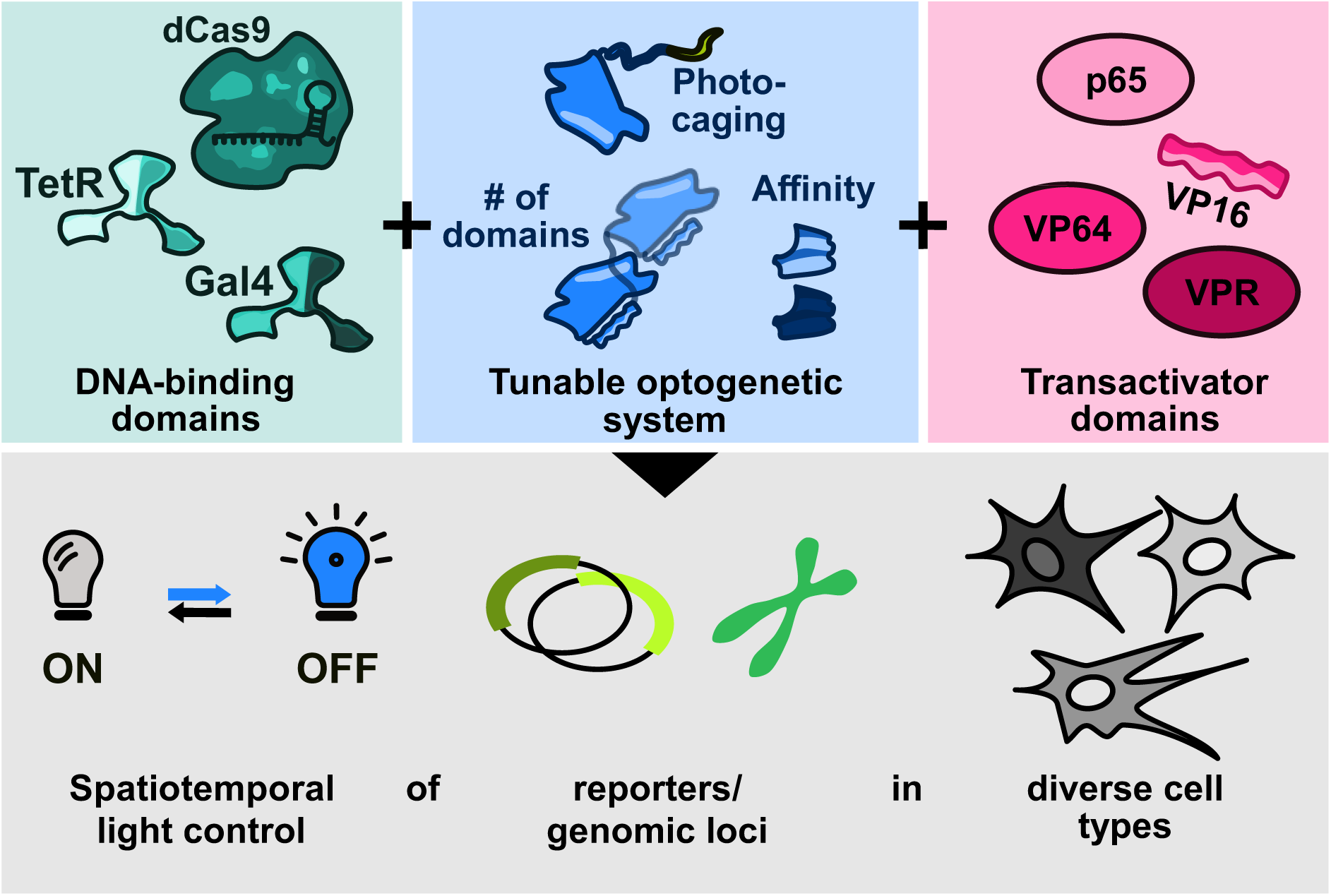
LOOMINA – a modular toolbox for the tunable optogenetic control of transcriptional induction. The combination of the LOVTRAP dissociation system with diverse DBDs and TADs enables the customized optogenetic transcriptional control of reporter expression or endogenous genes in various human cell lines.

LOOMINA, in contrast, is in itself a whole toolbox consisting of different components that can be readily combined and interchanged (**Fig. 5**). In addition to its modularity, we demonstrated LOOMINA’s robust functionality in a variety of settings, from reporter assays, to the control of endogenous genes, to spatial transcriptional regulation via a custom photomask. While related studies on Cas9-based optogenetic transcriptional activators have focused almost exclusively on HEK293T cells^3–5^, we have thoroughly characterized LOOMINA in various cell lines including HeLa, HepG2, U2OS and N2a.

The dynamic range of LOOMINA can be dramatically altered by varying the DBD:TAD vector mass ratio (**Fig. 1g-i**, **Supplementary Fig. 6**). Providing the vector encoding the much larger Cas9 transgene in excess has been proven beneficial in all settings. Unlike most two-hybrid systems that operate in a 1:1 stoichiometry, LOOMINA incorporates three *As*LOV2 domains, each of which can bind one Zdk-TAD protein. Mechanistically, the concatenation of more than one LOV2 domain is a straightforward way to improve the performance of an optogenetic switch. While we are not aware of any other reports using this strategy, it may be easily transferable to other LOV2-based dimerization systems, such as iLID^29^.

Finally, LOOMINA fills a gap in the optogenetic arsenal for transcriptional control by enabling researchers to turn off gene expression with blue light in a highly customizable manner. This mechanism of action further broadens the applicability of optogenetic transcriptional regulators and will be particularly important if transcriptional induction is to be reversed at a given time point in an experiment. With transcriptional induction being its default state in the dark, LOOMINA’s mode of action aligns well with various experimental setups and will enable researchers to reduce the required illumination time in many cases. In this sense, LOOMINA may even be a well-suited candidate for future optogenetic investigations *in vivo*, where continuous illumination tends to be more challenging and its duration is thus limited.

In conclusion, LOOMINA represents a powerful toolkit for CRISPR-mediated light off-operated transcriptional control that can be used in various configurations and experimental setups and will be a valuable addition to the existing set of optogenetic strategies for gene regulation.

## Methods

### Construct design and cloning

All plasmids generated and used in this study are listed in **Supplementary Table 1**. The corresponding amino acid sequences of the relevant fusion proteins are shown in **Supplementary Table 2**. Annotated plasmid sequences are provided in the supplementary data as genbank files. Vectors were either constructed by standard restriction-ligation cloning or via Golden Gate Assembly^44^. In brief, custom DNA oligonucleotides and double-stranded DNA were ordered from IDT or Merck. DNA fragments required for cloning were PCR amplified by either Q5 Hot Start high-fidelity DNA polymerase (New England Biolabs), the PrimeSTAR Max DNA Polymerase (Takara Bio) or Platinum Superfi II DNA polymerase (Thermo Fisher Scientific) according to the manufacturer’s protocols. PCR Products were separated by agarose gel electrophoresis and the desired bands were cut out followed by extraction of the DNA using the QIAquick gel extraction kit (Qiagen) or the Nucleospin gel and PCR clean-up kit (Macherey-Nagel). Restriction and ligation reactions were performed using enzymes and buffers obtained from New England Biolabs and Thermo Fisher Scientific. Finally, chemically competent TOP10 *E. coli* cells (Thermo Fisher Scientific) or One Shot Stbl3 *E. coli* bacteria (Invitrogen) were transformed with the assembled constructs. Plasmid DNA was purified using the QIAprep Spin Miniprep, Plasmid Plus Midi kit (QIAGEN) or NucleoSpin Plasmid kit (Macherey-Nagel). The correct assembly of all plasmids was verified by Sanger sequencing (Microsynth Seqlab).

The dCas9-AsLOV2 constructs were obtained from the EF1a driven, mammalian expression dCas9-VP64 vector previously reported by our group^45^. The Tet-inducible reporter constructs and corresponding sgRNA expression plasmid were a kind gift from Moritoshi Sato (Addgene-IDs: #64161, #64127, #64128). The mCherry-P2a DNA sequence was amplified from Addgene plasmid #125148, which was a kind gift from Stanley Qi. The Zdk sequences were either amplified by PCR or ordered as dsDNA from IDT. The sgRNA sequences for the endogenous targets *IL1RN, MYOD* and *OCT4* were adapted from previous studies by Perez-Pinera et al.^46^ and Hu et al.^47^ and cloned into U6 driven sgRNA expression vector that was as a kind gift from Dirk Grimm. The sgRNAs targeting the promoter region of the murine genes *Arc, Kir2.1* and *Neurog2* were custom designed using CRISPOR online tool (http://crispor.tefor.net)^48^ and individually cloned into the plasmid pLenti-SpBsmBI-sgRNA-Hygro (Addgene-ID: #62205; a kind gift from Rene Maehr), containing an U6 driven sgRNA scaffold. For *Arc* and *Kir2.1,* 5 and 6 sgRNAs were multiplexed into the same plasmid, respectively, using NEBridge Golden Gate Assembly kit (NEB), with custom designed primers. A scrambled sgRNA plasmid targeting the bacterial *LacZ* gene, which is not present in the human or murine genome was constructed for negative controls.

### Cell culture

All cell lines used in this study were maintained in 1xDMEM without phenol red (Thermo Fisher Scientific) supplemented with 10% (v/v) bovine fetal calf serum (Thermo Fisher Scientific), 2 mM L-glutamine, 100 U/mL penicillin and 100 μg/mL streptomycin (all Thermo Fisher Scientific). Cells were incubated at 37 °C and 5 % CO2 in a humidified incubator and confluence was maintained between 70 %-90 % by regular passaging. HEK293T and HeLa cells were received as a kind gift by Dirk Grimm, Heidelberg University Hospital; U2OS cells obtained from Barbara Di Ventura, Albert-Ludwigs-University Freiburg; HepG2 cells were a kind gift from Stephan Herzig, Helmholtz Diabetes Center Munich. Neuro-2a (N2a) cells were supplied by ATCC and banked at EPFL. Prior to use, the cell lines were authenticated and tested for mycoplasma contamination.

### Blue light setup

To achieve homogeneous blue light illumination, a custom blue light setup was used that consisted of six blue light high-power LEDs (type CREE XP-E D5–15; emission peak ∼460 nm; emission angle ∼130°; LED-TECH.DE) connected to a Switching Mode Power Supply (Manson; HCS-3102). The LEDs were controlled by Raspberry Pi with a custom python script. Cell culture dishes and well-plates were illuminated from underneath with a light intensity of ∼5 W/m^2^, which was regularly controlled with a LI-COR LI-250A light meter. To avoid phototoxic effects a duty cycle of 5 s blue light illumination followed by 10 s in darkness was continuously executed during all illumination periods. The illuminated and dark samples were kept in the same incubator during exposure, while the dark samples were shielded with a black vinyl foil (Starlab) and stored in a closed nontransparent box.

Experiments in N2a cells were performed by in another lab using a different blue light setup. In brief, light exposure was achieved using a LED Array Driver LAD-1 (Amuza Inc., #E55.100.01). In order to avoid heating and to be able to test different light intensities, one or more dark plexiglass films were placed on top of the LED Array plate. The LED Array plate was positioned inside a cell culture incubator, set on TRG mode and connected to a Master-8-vp (A.M.P. Instruments LTD.), which allowed to control light duty cycles as described in the figures. Light intensity was controlled using a Thorhlabs Power Meter PM100D light meter and kept at 200 Lux ≘ 4.9 W/m^2^. This light intensity corresponded to a voltage intensity of 7.52 V. Cells were exposed to blue-light illumination for 48 hours with illumination intervals of 1 s light-on and 1 s light-off, unless otherwise stated. The dark controls were kept in the same incubator, but covered by aluminum foil.

### Luciferase reporter assay

HEK293T cells were seeded at a density of 12,500 cells per well in a 96-well plate. HeLa, U2OS and HepG2 cells were seeded at a density of 6,000 cells per well. 24 h after seeding, the cells were transfected using Lipofectamine 3000 (Thermo Fisher Scientific) according to the manufacturer’s protocol with 0.3 μL Lipofectamine per well. A total of 100 ng DNA per well was used in all experiments. For the Gal4-and TetR-based luciferase reporter assays, 20 ng of the pRL-TK RLuc construct and 20 ng of the Tet-inducible FLuc reporter were transfected together with 40 ng of the respective DBD-TAD control plasmid or plasmids expressing the LOOMINA components in the indicated ratios. The total amount of DNA was topped up to 100 ng using an empty pBluescript stuffer plasmid. For the initial dCas9 reporter assays (Supplementary figures 4-6), an sgRNA expressing plasmid was co-transfected as an additional component. For all following experiments a construct that co-expresses the *Renilla* luciferase and a respective sgRNA was used instead. 40 ng of either the CRISPRa control constructs or both dCas9-AsLOV2 and Zdk3-TAD were used. The respective vector mass ratios are indicated in the figures. Finally, the DNA was topped up to 100 ng using an empty pBluescript stuffer plasmid. 3 h after transfection, one sample was placed under blue light exposure. After 48 h, the medium was removed and the samples were lysed in Passive Lysis buffer (Promega) for 40 min in the case of HEK293T cells and 1 h in the case of the other cell lines used. Finally, luciferase activity was assessed in a microplate reader (M Plex plate reader, Tecan) using the Dual Glo Luciferase Assay System (Promega). Luciferase substrate injection was followed by a 2 s settling time and 8 s of luminescence signal integration. Finally, the firefly luciferase photon counts were normalized to the *Renilla* luciferase photon counts, which served as internal expression control.

For luciferase experiments with N2a cells, the same protocol was followed with minor modifications. N2a cells were seeded at a density of 12,500 cells per well. The same amounts of DNA were transfected using Lipofectamine 2000 instead of Lipofectamine 3000, according to the manufacturer’s instructions. pCMV6-empty was used as a stuffer plasmid. 8 hours post-transfection, the medium was changed to DMEM and one plate was set under blue light exposure. For luciferase measurements, an integration time of only 1 s was used for N2a cells

### RT-qPCR

For RT-qPCR experiments, HEK293T cells were seeded in 12 well plates at a density of 1x10^5^ cells per well, HeLa cells were seeded with 0.6x10^5^ cells per well. The next day, cells were transfected using Lipofectamine 3000 (Thermo Fisher Scientific) according to the manufacturer’s protocol. A Lipofectamine concentration of 1,5 μL/well was chosen. A total of 1000 ng of plasmid DNA was transfected per well, comprising 75 ng of each sgRNA expression plasmid targeting different positions in the GOI promoter region (**Supplementary Table 3**) and 400 ng of the CRISPRa control or dCas9-AsLOV2 and Zdk3-TAD constructs, respectively. The vector mass ratios between the dCas9 and TAD constructs are indicated in the figures. In some cases, 700 ng of the dCas9 and TAD constructs were transfected, as indicated in the figures. An empty pBluescript stuffer plasmid was used to adjust the amount of transfected DNA to 1,000 ng per well. Illumination of one replicate was started 3 hours post transfection, while the other replicate was permanently kept in darkness. Cells were lysed after 72 h using TRI reagent (Zymo Research). Total RNA was extracted using a Direct-zol RNA Microprep Kit (Zymo Research) and 1,000 ng was used as input for reverse transcription, primed by random oligos using the RevertAid RT Kit (Thermo Fisher Scientific). The PowerSYBR Green PCR Master Mix (Thermo Fisher Scientific) was used for qPCR analysis of the cDNA with 0.5 μM primer and 1 μL of the RT reaction as template. qPCR primers for *GAPDH*, *IL1RN* and *MYOD* were obtained from literature^45,46^ while *OCT4* primers were designed by us. All oligonucleotides used for RT-qPCR are listed in **Supplementary table 4**. RT-qPCR was performed on a Q qPCR cycler (Quantabio) using the following program: 10 min at 95 °C for initial denaturation followed by 45-50 cycles of 95 °C for 15 s and 60 °C for 60 s with a final melting curve of 60-95 °C. CT values were assigned based on identical thresholds between all replicates of one measurement. Gene expression levels were calculated by the ΔΔC_T_-method^49^ with *GAPDH* as the constitutive control gene.

RT-qPCR experiments with N2a cells were performed in a different lab under the following conditions: For RT-qPCR, the day prior to transfection N2a cells were seeded in two 12-well plates with a density of 350,000 cells per well. A total amount of 1,000 ng DNA per well was transfected in Opti-MEM reduced serum medium with 2% Lipofectamine 2000 according to the manufacturer’s instruction. For the CRISPRa control, an equimolar quantity of dCas9-TAD vector, sgRNA vector and pCMV6-empty stuffer plasmid were used. The vector mass ratios between the dCas9-AsLOV2 and Zdk3-TAD constructs are, instead, indicated in the figures. In the latter case, the quantity of sgRNA vector used was in equimolar ratio to the dCas9-AsLOV2 vector. Eight hours post-transfection, the medium was changed to DMEM and one plate was set under blue light exposure.

Cells were lysed using Buffer RLT (Qiagen) and homogenized using either syringes and needles or QIAshredder (Qiagen). RNA was then extracted using QIAGEN RNeasy Mini Kit (Qiagen), following manufacturer’s instructions. RNA was eluted in 30 μL of RNAse-free water and quantified using a Nanodrop 1000 spectrophotometer. Then, 2 μg of RNA were reverse-transcribed into cDNA using SuperScript III First-Strand Synthesis (Invitrogen). Samples were cycled in a T100 Thermal Cycler (Bio-Rad), according to the protocol supplied with the synthesis kit, and the parental RNA strand was removed using RNAse H (Invitrogen). Each sample was then diluted 1:5 using RNAse-free water. For RT-qPCR, a mix of 2.7 μl water, 5 μl of SYBR Green Mastermix (Applied Biosystems), 0.3 μl of the forward and reverse intron-spanning primers at 5 μM was prepared for each sample. All primers were designed in-house and are listed in Supplementary table 4. 8 μl of the mix and 2 μl of cDNA was pipetted in a MicroAmp Fast Optical 96-well reaction plate (Applied Biosystems) in triplicates. The RT-qPCR was carried out using a StepOnePlus Real-Time PCR System (Applied Biosystems). Reactions were subjected to 45-50 cycles of 95 °C for 15 s and 60 °C for 60 s. Gene expression levels were calculated with the ΔΔC_T_-method, using *Actin* as a constitutive control gene.

### Fluorescence Reporter Assay

For the photomask experiments in 35 mm culture dishes, 8x10^5^ cells were seeded 24 h before transfection. Cells were transfected using Lipofectamine 3000 (Thermo Fisher Scientific) with 5 μL p3000 reagent and 6.25 μL Lipofectamine 3000. A total of 1,500 ng plasmid DNA was transfected, consisting of 300 ng of the Tet-inducible mCherry reporter, 450 ng of the TetO-targeting sgRNA expression plasmid, 712.5 ng of the dCas9-AsLOV vector and 37.5 ng of the Zdk-TAD construct. 3 h post transfection a photomask was applied to the culture dishes using black vinyl foil (Starlab) and cells were incubated under blue light exposure. After 24 hours, the dishes were imaged using a Keyence BZ-9000 microscope. Multiple images were taken and stitched using the microscopes software. The fluorescence intensity was quantified along the x-axis of the photo via ImageJ (Version 1.54).

### RNAscope experiment

Cells were plated on two Millicell EZ slides (Merck) at a density of 150,000 cells per well and transfected with 400 ng of plasmid DNA per well. The vector mass ratio between the dCas9-AsLOV2 and Zdk3-TAD constructs was 3:1, while the sgRNAs vector was equimolar to the dCas9 construct. Transfection was performed using Opti-MEM (Thermofisher). Media was changed to DMEM with GlutaMAX after 8 hours and one slide was illuminated on the LED Array for 48 hours, while the other was kept in darkness.

RNA fluorescent in situ hybridization (RNAfish) experiments were carried out using RNAscope Multiplex Fluorescent Kit v2 (Bio-techne), following manufacturer’s instructions for formalin-fixed samples. Briefly, chambers were dismantled and cells were washed with PBS and fixed in 10% Neutral Buffered Formalin (NBF). Next, cells were washed with PBS, dehydrated with an increasing concentration of EtOH and stored at -20°C until the next step. After rehydration, RNAscope Hydrogen Peroxide was added, cells were washed and an Immedge Hydrophobic barrier was drawn around each well. The cells were then treated with RNAscope Protease III. The Mm Arc-C2 probe (Bio-Techne) was hybridized for 2 hours at 40 °C. The signal of the probe was then amplified three times, using RNAscope Multiplex FL v2 Amp, and incubated with TSA Opal650 (Akoya Biosciences) diluted 1/1500. After 30 minutes blocking with 1% BSA, slides were incubated overnight at 4°C with primary antibodies: goat anti GFP (1:400, Abcam), rabbit anti RFP (1:500, Abcam) and mouse anti Cas9 (1:500, Active Motif). After incubation with secondary antibodies (donkey anti goat Alexa 488 (1/800, Thermo Fisher), donkey anti rabbit Alexa 568 (Thermo Fisher) and donkey anti mouse Alexa 750 (Abcam), cells were counterstained with DAPI and mounted with Prolong Gold Antifade Mounting medium (Thermo Fisher). Slides were imaged by a Zeiss confocal LSM980, using a 63x-oil objective with tile and doing an orthogonal projection of a z-stack of at least 12 layers, 0.7 μm wide each. For quantification, the same slides were imaged using the Olympus Slide Scanner VS200, with a 20x objective.

### Image analysis and quantification

For dCas9, GFP and mCherry triple-positive cell detection, images acquired with a Slide Scanner VS200 were analyzed using QuPath 0.4.3. Briefly, nuclei were automatically assigned via the DAPI signal and each nucleus expanded by 5 μm, to be considered as a “cell”. Triple-positive cells were identified by an automatically set threshold for each of the three channels and confirmed manually. Data was exported from QuPath and analyzed using a custom-made Python script to determine the percentage of triple-positive cells in each image. The DAPI signal was uniform across different images.

### Statistical analysis

Biological replicates refer to individual experiments with cells seeded, transfected and analyzed on different days. Technical replicates were created for luciferase and fluorescence reporter assays by transfecting individual wells of a microtiter plate on the same day. Data points represent individual replicates, as indicated in the figure legends. Bars represent the means of replicates and the error bars indicate the standard deviation. Fold changes are indicated, if relevant. Statistical analysis was performed with unpaired Student t-test or two-way ANOVA analysis of variance. A p-value below 0.05 was considered significant (marked by asterisks in the graphs). Data analysis was performed using the GraphPad Prism 10 software (GraphPad Software) or custom Python (3.8) scripts.

## Data Availability

Plasmid maps of the lead candidates are provided as genbank files. Lead constructs for pcVP16 and LOOMINA will also be made available via Addgene. Otherwise, all data is contained within the figures and will be made available upon reasonable request.

## Acknowledgements

We thank the members of the Niopek lab for helpful discussions. We also thank the workshop at the Biology Department of the Technical University Darmstadt for the construction of customized illumination setups.

## Author contributions

D.N., B.C., J.G. and J.M. conceived the study. P.M., N.S., M.F., D.C. and S.A. designed and performed the experiments. P.M., N.S., M.F. and J.M. analyzed the data. D.N. and J.M. directed the work. D.N. secured funding. J.M., P.M., M.F. and D.N. wrote the manuscript with support from all authors.

## Additional information

The online version contains supplementary material available at xxxxx.

