## Supplementary Information for "A modular toolbox for the optogenetic deactivation of transcription"

### Content:

Supplementary Fig. 1-13

Supplementary Table 1-4

Supplementary References

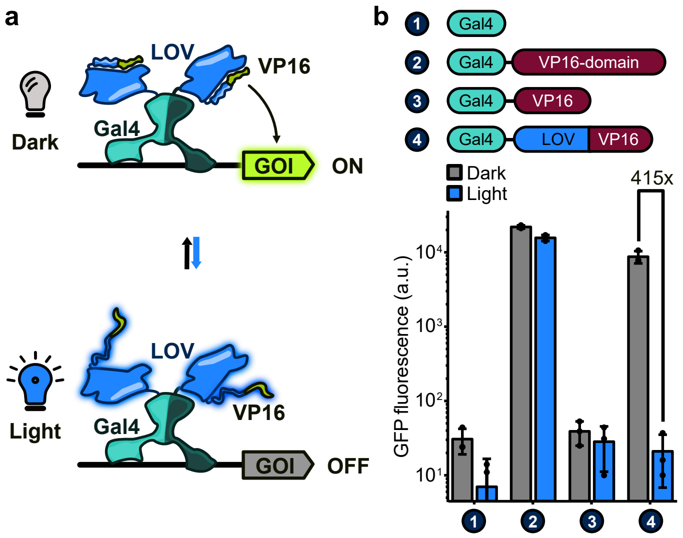


**Supplementary Fig. 1: The VP16 peptide potently activates transcription only in the *As*LOV2 context.**

**a** Schematic representation of the proposed VP16-photocaging mechanism. GOI: gene of interest. **b** Initial assessment of pcVP16 in comparison to the full VP16 domain and the VP16 transactivating peptide. HEK293T cells were transfected with plasmids expressing (i) a Gal4-UAS-controlled GFP reporter and (ii) the corresponding Gal4 construct, as indicated by the circled numbers. Following 20 h of incubation under blue-light or in the dark, GFP expression was quantified in a plate reader. Bars represent means of three technical replicates. Error bars indicate the S.D. Individual data points are marked as dots. Fold changes between samples incubated with or without blue light exposure are shown for the optogenetic variants.

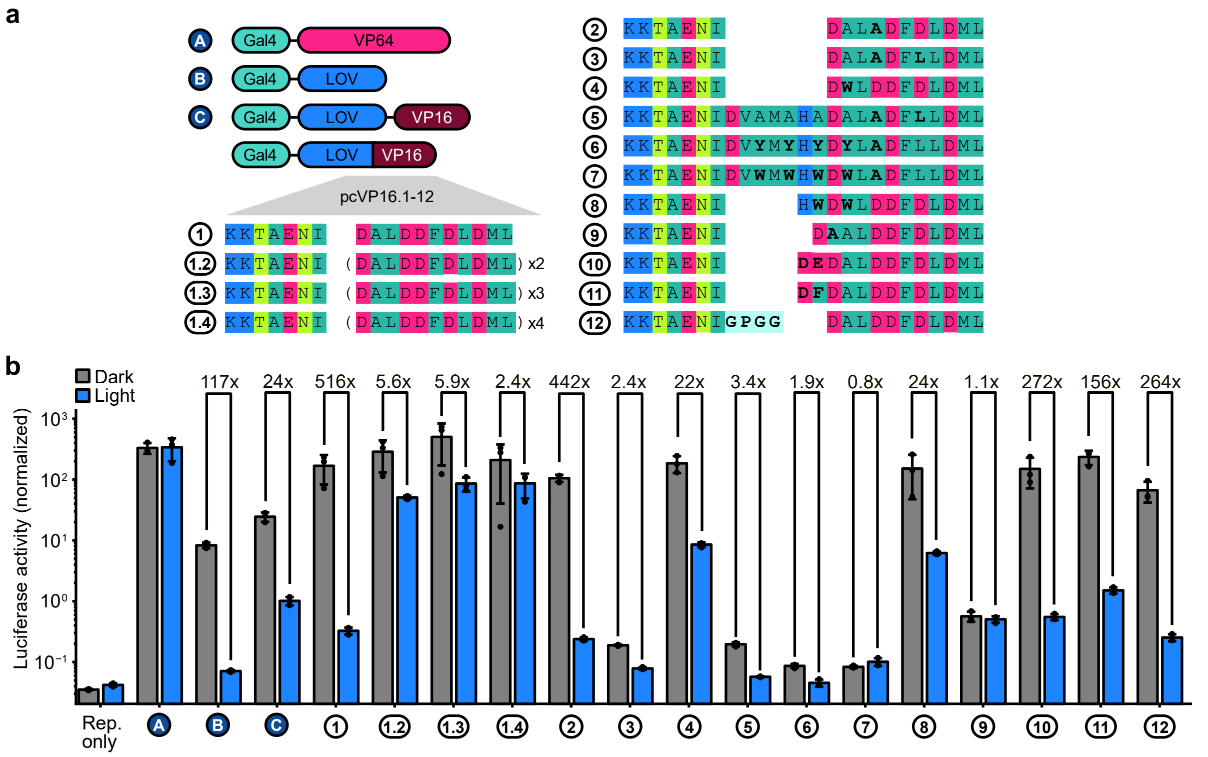


**Supplementary Fig. 2: Screening and optimization of light-switchable pcVP16 transcriptional activators.** **a** Domain architectures and amino acid sequences of the photocaged hybrid peptides are depicted. **b** Screening of different pcVP16-64 designs. HEK293T cells were transfected with plasmids encoding (i) a Gal4-inducible firefly luciferase, (ii) a *Renilla* luciferase and (iii) the Gal4-pcVP16.1-12 variants from **a**, as indicated by the circled numbers. Samples were incubated under blue light exposure or in the dark and luciferase activity was measured after 48 h. Bars represent means of three technical replicates. Error bars indicate S.D. Individual data points are marked as dots. Fold changes between samples incubated with or without blue light exposure are shown. Rep. only: reporter only control.

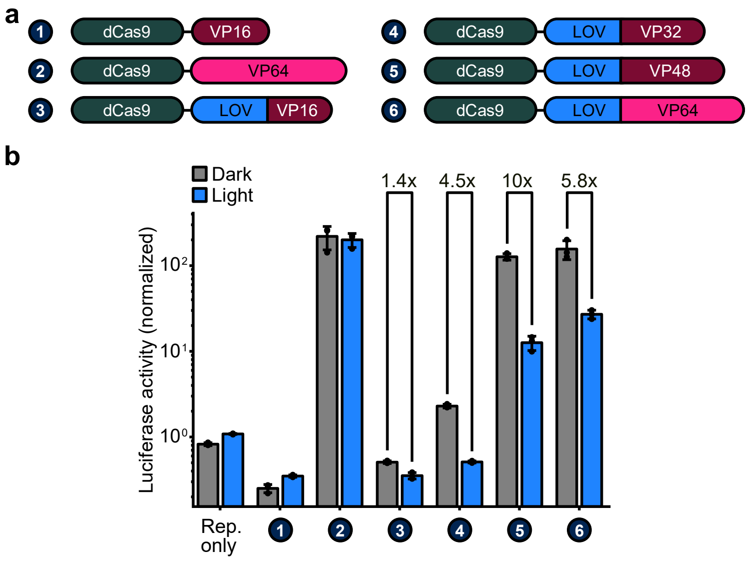


**Supplementary Fig. 3: The VP16 reverse photocaging strategy is compatible with dCas9 as DNA-binding domain.**

**a** Domain architectures of the tested pcVP16-64 designs in combination with dCas9 as programmable DBD. **b** Evaluation of pcVP16-64 in combination with dCas9. HEK293T cells were transfected with plasmids encoding (i) a Tet-inducible luciferase, (ii) a *Renilla* luciferase, (iii) a Tet-targeting sgRNA and (iv) the indicated dCas9-VP16-64 construct. Samples were incubated under blue light exposure or in the dark and luciferase activity was measured after 48 h. Bars represent means of three technical replicates. Error bars indicate S.D.. Individual data points are marked as dots. Fold changes between samples incubated with or without blue light exposure are shown. Rep. only, reporter only control. Rep. only: reporter only control.

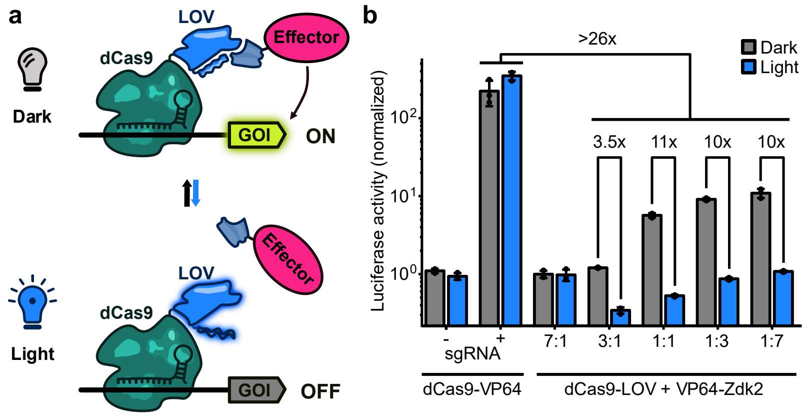


**Supplementary Fig. 4: A two-hybrid transactivation approach using a dCas9-LOV fusion in combination with a C-terminal fusion of Zd2k to VP64 enables modest transcriptional activation.**

**a** Schematic mechanism of light-controllable transcriptional activation with dCas9-LOVTRAP. GOI: gene of interest. **b** HEK293T cells were transfected with plasmids containing (i) a Tet-inducible luciferase reporter, (ii) a constitutively active *Renilla* luciferase, (iii) a TetO-targeting sgRNA and (iv) components of the optogenetic dCas9/TAD system or respective controls, as indicated in the figure. The dCas9:TAD vector mass ratios are shown below the bars. Samples were incubated under blue light or in the dark and luciferase activity was measured after 48 h. Bars represent the mean of three technical replicates. Error bars indicate the S.D. and data points the individual replicates. Fold changes between illuminated and dark samples are indicated.

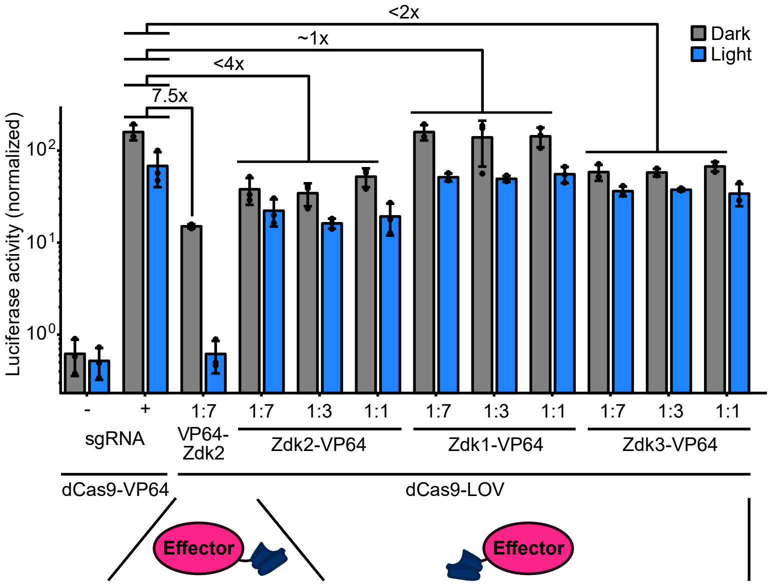


**Supplementary Fig. 5: N-terminal fusions of Zdk to the effector domain drastically enhance transcriptional activation.**

HEK293T cells were transfected with plasmids encoding (i) a TetO-regulated firefly luciferase, (ii) a *Renilla* luciferase, (iii) a TetO-targeting sgRNA and (iv) the indicated optogenetic dCas9/TAD combinations or controls. The dCas9:TAD vector mass ratios are shown below the bars. Samples were illuminated with blue light or kept in the dark for 48 h, followed by a luciferase assay. Bars represent the mean of three technical replicates. Error bars indicate the S.D. and data points the individual replicates. Fold changes between the positive control and individual transactivator configurations are indicated.

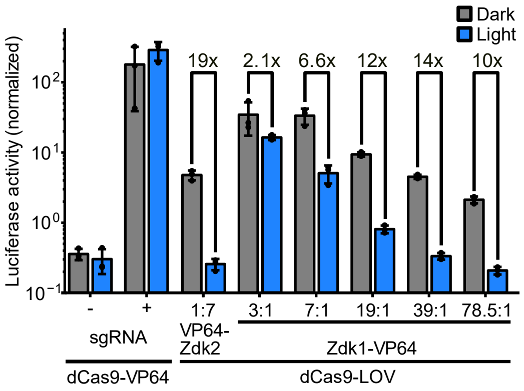


**Supplementary Fig. 6: Tuning construct stoichiometry enables controlling the system's dynamic range of gene regulation and overall activation.**

HEK293T cells were transfected with plasmids encoding (i) a TetO-regulated firefly luciferase, (ii) a *Renilla* luciferase, (iii) a TetO-targeting sgRNA and (iv) the indicated optogenetic dCas9/TAD combinations or controls. The dCas9:TAD vector mass ratios are shown below the bars. Samples were illuminated with blue light or kept in the dark for 48 h, followed by a luciferase assay. Bars represent the mean of three technical replicates. Error bars indicate the S.D. and data points the individual replicates. Fold changes between illuminated and dark samples are indicated.

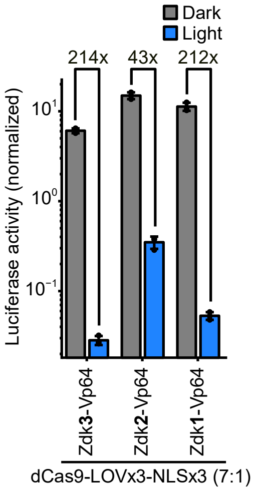


**Supplementary Fig. 7: The choice of the specific Zdk affects the systems leakiness and dynamic range.** HEK293T cells were transfected with plasmids encoding (i) a TetO-regulated firefly luciferase, (ii) a *Renilla* luciferase together with a TetO-targeting sgRNA and (iii) dCas9-AsLOVx3-NLSx3 and (iv) a Zdk-VP64 variant using a 7:1 plasmid mass ratio. Samples were illuminated with blue light or kept in the dark for 48 h, followed by a luciferase assay. Bars represent the mean of three biological replicates, each consisting of three technical replicates. Error bars indicate the S.D. and data points the individual values of biological replicates. Fold changes between illuminated and dark samples are indicated.

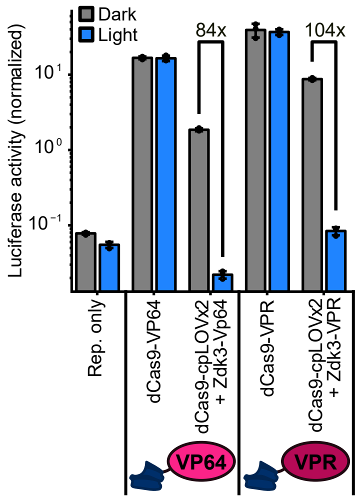


**Supplementary Fig. 8: LOOMINA can be used in conjunction with cpLOV as light-sensing domain.**

HEK293T cells were transfected with plasmids encoding (i) a TetO-controlled firefly reporter, (ii) a constitutive *Renilla* luciferase together with a TetO-targeting sgRNA and (iii) dCas9-cpAsLOV2x2 with either (iv) Zdk3-VP64 or Zdk3-VPR in a 3:1 or 7:1 vector mass ratio, respectively. Fusions of dCas9 to VP64 or VPR were transfected in the control samples. Samples were illuminated with blue light or kept in the dark for 48 h, followed by a luciferase assay. Bars represent the mean of three biological replicates, each consisting of three technical replicates. Error bars indicate the S.D. and data points the individual values of biological replicates. Fold changes between illuminated and dark samples are indicated. Rep. only, reporter only control.

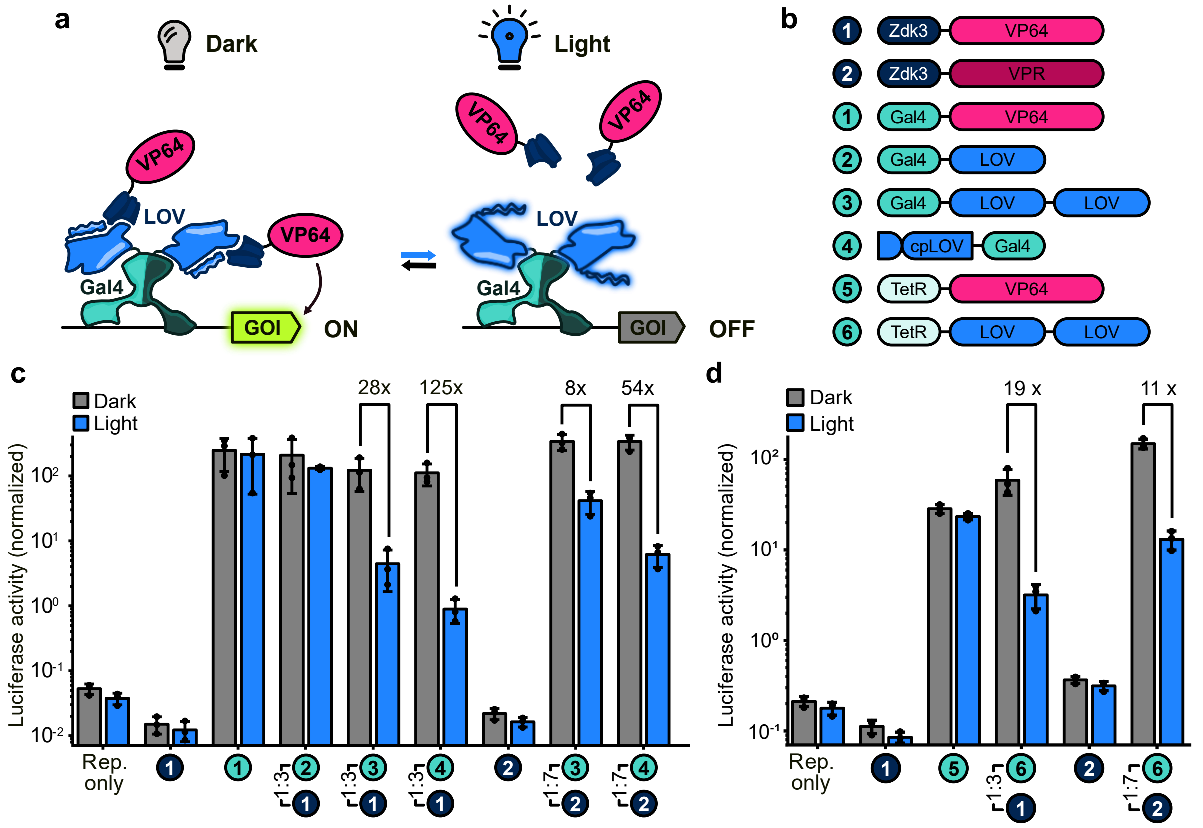


**Supplementary Fig. 9: LOOMINA is compatible with different DBDs in a plug-and-play manner.**

**a** Schematic representation of the Gal4-LOVTRAP transcription control circuit. GOI: gene of interest. **b** Domain architectures of the optogenetic Gal4/TetR-TADs. **c**, **d** HEK293T cells were transfected with plasmids encoding (i) a Gal4- (**c**) or Tet-inducible (**d**) luciferase reporter, (ii) a constitutively active *Renilla* luciferase and (iii) the respective Gal4- (**c**) or TetR-TAD (**d**) combinations, as indicated by the circled numbers. A DBD:TAD vector mass ratio of 3:1 for VP64 or 7:1 for VPR was chosen. Samples were illuminated with blue light or kept in the dark for 48 h, followed by a luciferase assay. Bars represent the mean of three biological replicates. Error bars indicate the S.D. and data points the individual values from biological replicates. Fold changes between illuminated and dark samples are indicated. Rep. only: reporter only control.

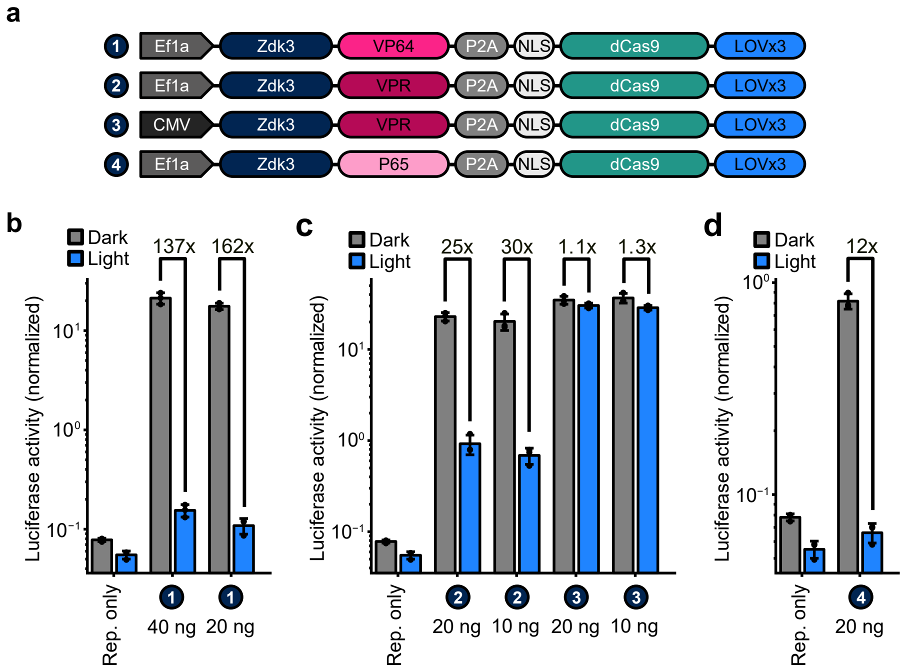


**Supplementary Fig. 10: The protein components of LOOMINA can be expressed from a single plasmid via P2A sequences.**

**a** Construct architectures of the P2A-separated single-transgene configurations of LOOMINA. **b-d** Dual-luciferase assays of HEK293T cells transfected with constructs encoding (i) a TetO-controlled luciferase reporter, (ii) a constitutively expressed *Renilla* luciferase together with a TetO-targeting sgRNA and (iii) the indicated amount of the respective P2A dCas9-LOVTRAP with either VP64 (**b**), VPR (**c**) or p65 (**d**) as TAD.Samples were illuminated with blue light or kept in the dark for 48 h, followed by a luciferase assay. Bars represent the mean of three biological replicates, each consisting of three technical replicates. Error bars indicate the S.D. and data points the individual values of the biological replicates. Fold changes between illuminated and dark samples are indicated. Rep. only, reporter only control.

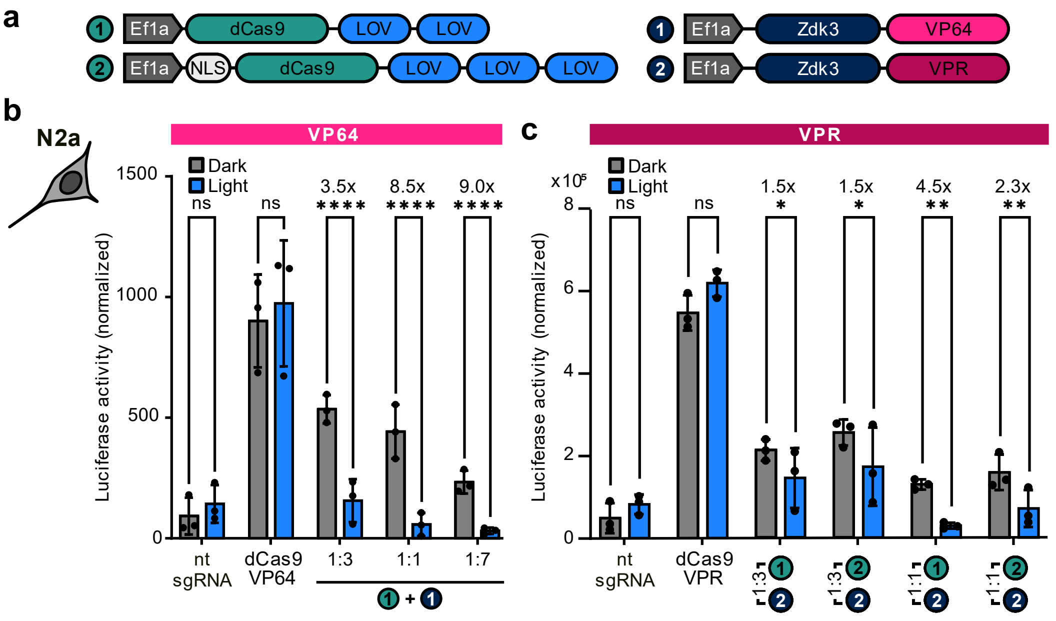


**Supplementary Fig. 11: LOOMINA enables light-controlled expression of a luciferase reporter in N2a cells.** **a** List of the used constructs. **b-c** N2a cells were transfected with plasmids encoding (i) a firefly luciferase downstream of 13x TetO sites, (ii) a *Renilla* luciferase and (iii) the indicated dCas9 and TAD components. As positive control, a direct dCas9-VPR fusion was used. The samples were either illuminated with a 5 s on/5 s off duty cycle or kept in darkness for 48 h, followed by luciferase measurements. Vector mass ratios between the dCas9-expressing construct and the transactivator construct are indicated. Bars represent means of three biological replicates. Data points show the individual biological replicates, which are the mean of three technical replicates. Error bars indicate S.D. Fold changes between illuminated and dark samples are indicated. Statistical significance was assessed by two-way ANOVA, with Sidak-correction for multiple comparison; n.s.: not significant; *p-value < 0.05; **p-value < 0.01, ****p-value<0.0001. Nt sgRNA: non-targeting sgRNA control.

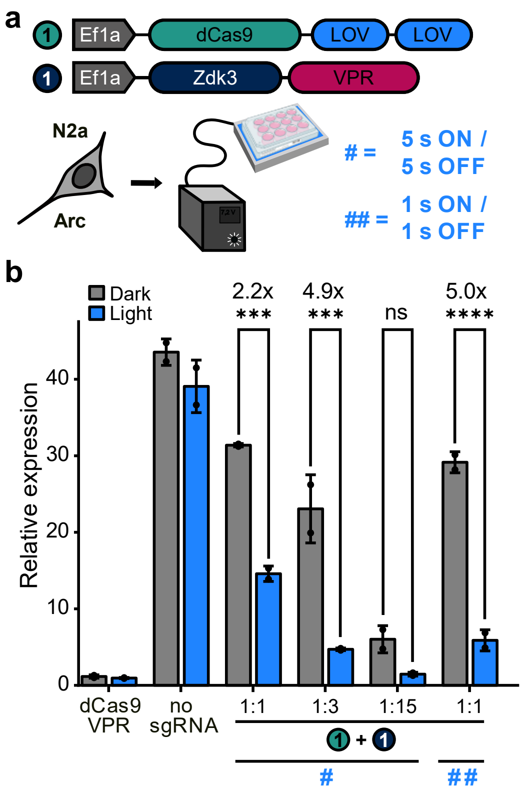


**Supplementary Fig. 12: Short illumination intervals can increase the performance of LOOMINA in N2a cells.** **a** List of the constructs used in the experiment. The tested illumination duty cycles are indicated. **b** N2a cells were transfected with the indicated LOOMINA components alongside constructs encoding sgRNAs that target either the mouse gene *Arc* or non-targeting controls. Different illumination patterns were tested as indicated. As positive control, a direct dCas9-VPR fusion was used. The samples were either illuminated with the indicated on/off times for 48 hours or kept in darkness, followed by RT-qPCR analysis of gene expression. Bars represent means of two biological replicates. The vector mass ratios between the dCas9-expressing construct and the transactivator construct are indicated. Data points show the individual biological replicates, which are the mean of three technical replicates. Error bars represent the S.D. Fold changes between illuminated and dark samples are indicated. Statistical significance was assessed by two-way ANOVA, with Sidak-correction for multiple comparison; n.s.: not significant; ***p-value < 0.001; ****p-value<0.0001. Nt sgRNA: non-targeting sgRNA control.

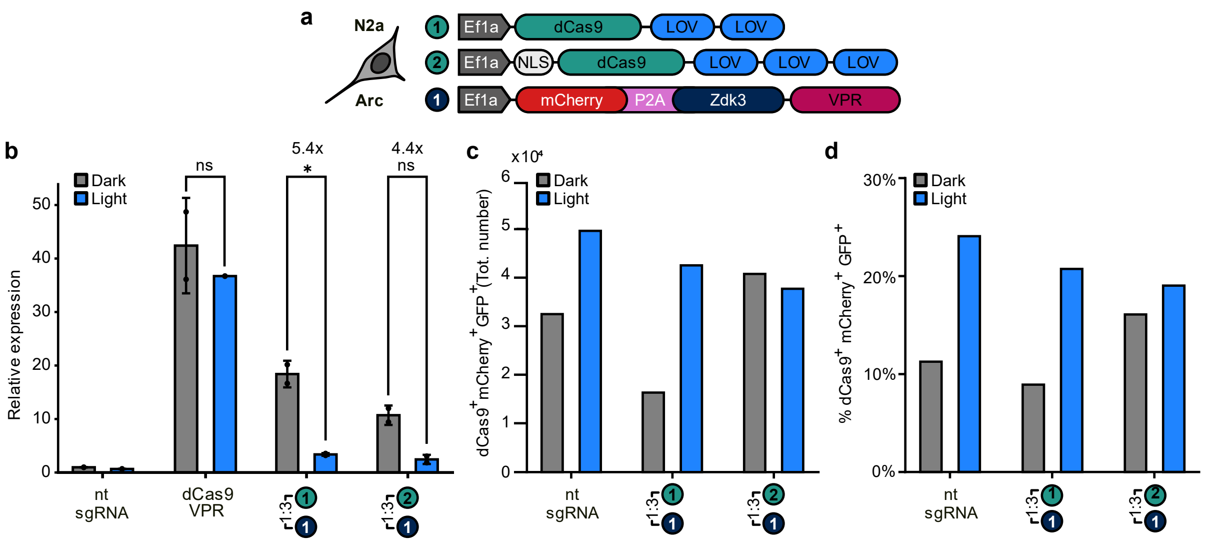


**Supplementary Fig. 13: RNAscope experiments reveal that 10-20% of the transfected N2a cells express all LOOMINA components.**

**a** Schematic of the used constructs. **b** N2a cells were transfected with constructs encoding the indicated LOOMINA components alongside plasmids expressing sgRNAs that target *Arc* or non-targeting controls. As positive control, a direct dCas9-VPR fusion was used. The samples were either illuminated for 48 hours or kept in darkness, followed by RT-qPCR analysis of gene expression. Bars represent means of two biological replicates. Data points show the individual biological replicates, which are the mean of three technical replicates. Error bars indicate S.D. Statistical significance was assessed by two-way ANOVA, with Sidak-correction for multiple comparison; n.s.: not significant; *p-value < 0.05. n = 2 replicates, except sample dCas9-VPR (light), which is n = 1 (one sample was lost during sample measurement). **c** Total number of triple positive cells detected in whole-slide RNAscope images. These cells were used for quantification of the *Arc* mRNA-probe displayed in **Fig. 4b-c**. **d** Percentage of triple positive cells over the total amount of cells detected in whole-slide images. **b-d** The circled numbers refer to the combination of Cas9 and TAD that was used. The vector mass ratios between the dCas9-expressing construct and the transactivator construct are indicated. Nt sgRNA: non-targeting sgRNA control.

**Supplementary Table 1: List of constructs used in this study.** UAS, upstream activation sequence; GFP: green fluorescent protein; Fluc: firefly luciferase; pc: photocaged; CMV: Cytomegaloviru; NLS: nuclear localization signal.

| # | Plasmid Name | Insert | Reference |
|---|---|---|---|
| 1 | pRL-TK | pTK; Renilla luciferase | Promega |
| 2 | pBluescript | Empty vector | Invitrogen |
| 3 | Gal4-UAS-GFP | Gal4-UAS; GFP reporter | This Study |
| 4 | Gal4-UAS-Fluc | Gal4-UAS; Firefly luciferase reporter | This Study |
| 5 | Tet-inducible Luciferase reporter | TetO; Luciferase reporter | [1] |
| 6 | Tet-inducible mCherry reporter | TetO; mCherry reporter | [1] |
| 7 | mCherry-MODC-Tet-ind. | TetO; mCherry-MODC | This Study |
| 8 | sgRNA TetO | U6; sgRNA against TetO | [1] |
| 9 | sgRNA-Tet-RLuc | U6; sgRNA against TetO. TetO-controlled Renilla luciferase | This Study |
| 10 | sgRNA IL1RN A | U6; sgRNA 1 against human Il1RN promoter | [2] |
| 11 | sgRNA IL1RN B | U6; sgRNA 2 against human Il1RN promoter | [2] |
| 12 | sgRNA IL1RN C | U6; sgRNA 3 against human Il1RN promoter | [2] |
| 13 | sgRNA IL1RN D | U6; sgRNA 4 against human Il1RN promoter | [2] |
| 14 | sgRNA MYOD A | U6; sgRNA 1 against human MYOD promoter | This Study |
| 15 | sgRNA MYOD B | U6; sgRNA 2 against human MYOD promoter | This Study |
| 16 | sgRNA MYOD C | U6; sgRNA 3 against human MYOD promoter | This Study |
| 17 | sgRNA MYOD D | U6; sgRNA 4 against human MYOD promoter | This Study |
| 18 | sgRNA OCT4 A | U6; sgRNA 1 against human OCT4 promoter | This Study |
| 19 | sgRNA OCT4 B | U6; sgRNA 2 against human OCT4 promoter | This Study |
| 20 | sgRNA OCT4 C | U6; sgRNA 3 against human OCT4 promoter | This Study |
| 21 | sgRNA OCT4 D | U6; sgRNA 4 against human OCT4 promoter | This Study |
| 22 | Gal4-VP16 full | CMV; SV40 NLS-Gal4-VP16 full domain | This Study |
| 23 | Gal4-VP16 | CMV; SV40 NLS-Gal4-VP16 | This Study |
| 24 | Gal4-VP64 | CMV; SV40 NLS-Gal4-VP64 | This Study |
| 25 | Gal4-VPR | CMV; SV40 NLS-Gal4-VPR | This Study |
| 26 | Gal4-pcVP16-1 | CMV; SV40 NLS-Gal4-AsLOV2-pcVP16-1 | This Study |
| 27 | Gal4-pcVP16-1.2 | CMV; SV40 NLS-Gal4-AsLOV2-pcVP16-1.2 | This Study |
| 28 | Gal4-pcVP16-1.3 | CMV; SV40 NLS-Gal4-AsLOV2-pcVP16-1.3 | This Study |
| 29 | Gal4-pcVP16-1.4 | CMV; SV40 NLS-Gal4-AsLOV2-pcVP16-1.4 | This Study |
| 30 | GAL4-pcVP16-2 | CMV; SV40 NLS-Gal4-AsLOV2-pcVP16-2 | This Study |
| 31 | GAL4-pcVP16-3 | CMV; SV40 NLS-Gal4-AsLOV2-pcVP16-3 | This Study |
| 32 | GAL4-pcVP16-4 | CMV; SV40 NLS-Gal4-AsLOV2-pcVP16-4 | This Study |
| 33 | GAL4-pcVP16-5 | CMV; SV40 NLS-Gal4-AsLOV2-pcVP16-5 | This Study |
| 34 | GAL4-pcVP16-6 | CMV; SV40 NLS-Gal4-AsLOV2-pcVP16-6 | This Study |
| 35 | GAL4-pcVP16-7 | CMV; SV40 NLS-Gal4-AsLOV2-pcVP16-7 | This Study |
| 36 | GAL4-pcVP16-8 | CMV; SV40 NLS-Gal4-AsLOV2-pcVP16-8 | This Study |
| 37 | GAL4-pcVP16-9 | CMV; SV40 NLS-Gal4-AsLOV2-pcVP16-9 | This Study |
| 38 | GAL4-pcVP16-10 | CMV; SV40 NLS-Gal4-AsLOV2-pcVP16-10 | This Study |
| 39 | GAL4-pcVP16-11 | CMV; SV40 NLS-Gal4-AsLOV2-pcVP16-11 | This Study |
| 40 | GAL4-pcVP16-12 | CMV; SV40 NLS-Gal4-AsLOV2-pcVP16-12 | This Study |
| 41 | Gal4-AsLOV2 | CMV; SV40 NLS-Gal4-AsLOV2 | This Study |
| 42 | Gal4-AsLOV2x2 | Ef1α; SV40 NLS-Gal4-AsLOV2x2 | This Study |
| 43 | cpLOV2-Gal4 | CMV; SV40 NLS-cpAsLOV2-Gal4 | This Study |
| 44 | TetR-VP64 | CMV; SV40 NLS-TetR-VP64 | This Study |
| 45 | TetR-AsLOV2x2 | CMV; SV40 NLS-TetR-AsLOV2x2 | This Study |
| 46 | dCas9-VP16 | Ef1α; SV40 NLS-dCas9-SV40 NLS-VP16 | This Study |
| 47 | dCas9-VP64 | Ef1α; SV40_NLS-dCas9-SV40_NLS-VP64 | [2] |
| 48 | dCas9-p65 | Ef1α; SV40 NLS-dCas9-SV40 NLS-p65 | This Study |
| 49 | dCas9-VPR | Ef1α; SV40 NLS-dCas9-SV40 NLS-VPR | This Study |
| 50 | dCas9-pcVP16 | Ef1α; SV40 NLS-dCas9-SV40 NLS-pcVP16 | This Study |
| 51 | dCas9-pcVP32 | Ef1α; SV40 NLS-dCas9-SV40 NLS-pcVP32 | This Study |
| 52 | dCas9-pcVP48 | Ef1α; SV40 NLS-dCas9-SV40 NLS-pcVP48 | This Study |
| 53 | dCas9-pcVP64 | Ef1α; SV40 NLS-dCas9-SV40 NLS-pcVP64 | This Study |
| 54 | dCas9-AsLOV2 | Ef1α; SV40 NLS-dCas9-AsLOV2 | This Study |
| 55 | dCas9-AsLOV2x2 | Ef1α; SV40 NLS-dCas9-AsLOV2x2 | This Study |
| 56 | dCas9-AsLOV2x3 | Ef1α; SV40 NLS-dCas9-AsLOV2x3 | This Study |
| 57 | dCas9-AsLOV2x3-NLSx3 | Ef1α; NLSx3-dCas9-AsLOV2x3 | This Study |
| 58 | CMV dCas9-AsLOVx3-NLSx3 | CMV; NLSx3-dCas9-AsLOV2x3 | This Study |

| | | | |
|---|---|---|---|
| 59 | dCas9-cpAsLOV2x2-Cterm | dCas9-cpAsLOV2x2 | This Study |
| 60 | Zdk2-VP64 | Ef1α; SV40 NLS-Zdk2-VP64-SV40 NLS | This Study |
| 61 | Zdk1-VP64 | Ef1α; SV40 NLS-Zdk1-VP64-SV40 NLS | This Study |
| 62 | Zdk3-VP64 | Ef1α; SV40 NLS-Zdk3-VP64-SV40 NLS | This Study |
| 63 | Zdk3-p65 | Ef1α; SV40 NLS-Zdk3-p65-SV40 NLS | This Study |
| 64 | Zdk1-VPR | Ef1α; SV40 NLS-Zdk1-VPR-SV40 NLS | This Study |
| 65 | Zdk3-VPR | Ef1α; SV40 NLS-Zdk3-VPR-SV40 NLS | This Study |
| 66 | CMV Zdk3-VPR | CMV; SV40 NLS-Zdk3-VPR-SV40 NLS | This Study |
| 67 | VP64-Zdk2 | Ef1α; SV40 NLS-VP64-Zdk2-SV40 NLS | This Study |
| 68 | Zdk3-VP64_P2A_dCas9-AsLOVx3-NLSx3 | Ef1α; SV40_NLS-Zdk3-VP64-SV40_NLS P2A NLSx3-dCas9-AsLOV2x3 | This Study |
| 69 | Zdk3-p65_P2A_dCas9-AsLOVx3-NLSx3 | Ef1α; SV40_NLS-Zdk3-p65-SV40_NLS P2A NLSx3-dCas9-AsLOV2x3 | This Study |
| 70 | Zdk3-VPR_P2A_dCas9-AsLOVx3-NLSx3 | Ef1α; SV40_NLS-Zdk3-VPR-SV40_NLS P2A NLSx3-dCas9-AsLOV2x3 | This Study |
| 71 | CMV_Zdk3-VPR_P2A_dCas9-AsLOVx3-NLSx3 | CMV; SV40_NLS-Zdk3-VPR-SV40_NLS P2A NLSx3-dCas9-AsLOV2x3 | This Study |
| 71 | mCheery P2A Zdk3-VPR | Ef1α; mCherry P2A SV40 NLS-Zdk3-VPR-SV40 NLS | This Study |

**Supplementary Table 2: Amino acid sequences of the key fusion proteins used in this study.** The construct numbers refer to Supplementary table 1. Grey: NLS; dark green: DBD; light blue: AsLOV2; light green: VP16 peptide; dark blue: Zd; pink: TAD; black: linkers.

| # | Plasmid name | Amino acid sequence |
|---|---|---|
| 26 | Gal4-pcVP16-1 | MPKKKRKVGGSGGSKLLSSIEQACDICRLKKLKCSKEKPKCAKCLKNNWECRYSPKTKRSPLTRAHLTEVESRLERLEQLFLLIFPREDLDMILKMDSLQDIKALLTGLFVQDNVNKDAVTDRLASVETDMPLTLRQHRISATSSSEESSNKGQRQLTVSGGSGGGSGGSSLATTLERIEKNFVITDPRLPDNPIIFASDSFLQLTEYSREEILGRNCRFLQGPETDRATVRKIRDAIDNQTEVTVQLINYTKSGKKFWNLFHLQPMRDQKGDVQYFIGVQLDGTEHVRDAAEREGVMLIKKTAENIDALDDFDLDML |
| 42 | Gal4-AsLOV2x2 | MPKKKRKVGGSGGSKLLSSIEQACDICRLKKLKCSKEKPKCAKCLKNNWECRYSPKTKRSPLTRAHLTEVESRLERLEQLFLLIFPREDLDMILKMDSLQDIKALLTGLFVQDNVNKDAVTDRLASVETDMPLTLRQHRISATSSSEESSNKGQRQLTVSSAGGGGSGLATTLERIEKNFVITDPRLPDNPIIFASDSFLQLTEYSREEILGRNCRFLQGPETDRATVRKIRDAIDNQTEVTVQLINYTKSGKKFWNLFHLQPMRDQKGDVQYFIGVQLDGTEHVRDAAEREGVMLIKKTAENIDEAAKELGSGSGSGSGLATTLERIEKNFVITDPRLPDNPIIFASDSFLQLTEYSREEILGRNCRFLQGPETDRATVRKIRDAIDNQTEVTVQLINYTKSGKKFWNLFHLQPMRDQKGDVQYFIGVQLDGTEHVRDAAEREGVMLIKKTAENIDEAAKEL |
| 43 | cpLOV2-Gal4 | MTEHVRDAAEREGVMLIKKTAENIDEAAKELGGGSGGSGGGLATTLERIEKNFVITDPRLPDNPIIFASDSFLQLTEYSREEILGRNCRFLQGPETDRATVRKIRDAIDNQTEVTVQLINYTKSGKKFWNLFHLQPMRDQKGDVQYFIGVQLDGGGGGSPKKKRKVGGSGGSKLLSSIEQACDICRLKKLKCSKEKPKCAKCLKNNWECRYSPKTKRSPLTRAHLTEVESRLERLEQLFLLIFPREDLDMILKMDSLQDIKALLTGLFVQDNVNKDAVTDRLASVETDMPLTLRQHRISATSSSEESSNKGQRQLTVSGGSGGGSGGSS |
| 45 | TetR-AsLOV2x2 | MSPKKKRKVEASMTRLDKSKVINSALELLNEVGIEGLTTRKLAQKLGVEQPTLYWHVKNKRALLDALAIEMLDRHHTHFCPLEGESWQDFLRNNAKSFRCALLSHRDGAKVHLGTRPTEKQYETLENQLAFLCQQGFSLENALYALSAVGHFTLGCVLEDQEHQVAKEERETPTTDSMPPLLRQAIELFDHQGAEPAFLFGLELIICGLEKQLKCESGSSAGGGGSGLATTLERIEKNFVITDPRLPDNPIIFASDSFLQLTEYSREEILGRNCRFLQGPETDRATVRKIRDAIDNQTEVTVQLINYTKSGKKFWNLFHLQPMRDQKGDVQYFIGVQLDGTEHVRDAAEREGVMLIKKTAENIDEAAKELGSGSGSGSGLATTLERIEKNFVITDPRLPDNPIIFASDSFLQLTEYSREEILGRNCRFLQGPETDRATVRKIRDAIDNQTEVTVQLINYTKSGKKFWNLFHLQPMRDQKGDVQYFIGVQLDGTEHVRDAAEREGVMLIKKTAENIDEAAKEL |
| 54 | dCas9-AsLOV2 | MSPKKKRKVEASDKKYSIGLAIGTNSVGWAVITDEYKVPSKKFKVLGNTDRHSIKKNLIGALLFDSGETAEATRLKRTARRRYTRRKNRICYLQEIFSNEMAKVDDSFFHRLEESFLVEEDKKHERHPIFGNIVDEVAYHEKYPTIYHLRKKLVDSTDKADLRLIYLALAHMIKFRGHFLIEGDLNPDNSDVDKLFIQLVQTYNQLFEENPINASGVDAKAILSARLSKSRRLENLIAQLPGEKKNGLFGNLIALSLGLTPNFKSNFDLAEDAKLQLSKDTYDDDLDNLLAQIGDQYADLFLAAKNLSDAILLSDILRVNTEITKAPLSASMIKRYDEHHQDLTLLKALVRQQLPEKYKEIFFDQSKNGYAGYIDGGASQEEFYKFIKPILEKMDGTEELLVKLNREDLLRKQRTFDNGSIPHQIHLGELHAILRRQEDFYPFLKDNREKIEKILTFRIPYYVGPLARGNSRFAWMTRKSEETITPWNFEEVVDKGASAQSFIERMTNFDKNLPNEKVLPKHSLLYEYFTVYNELTKVKYVTEGMRKPAFLSGEQKKAIVDLLFKTNRKVTVKQLKEDYFKKIECFDSVEISGVEDRFNASLGTYHDLLKIIKDKDFLDNEENEDILEDIVLTLTLFEDREMIEERLKTYAHLFDDKVMKQLKRRRYTGWGRLSRKLINGIRDKQSGKTILDFLKSDGFANRNFMQLIHDDSLTFKEDIQKAQVSGQGDSLHEHIANLAGSPAIKKGILQTVKVVDELVKVMGRHKPENIVIEMARENQTTQKGQKNSRERMKRIEEGIKELGSQILKEHPVENTQLQNEKLYLYYLQNGRDMYVDQELDINRLSDYDVDAIVPQSFLKDDSIDNKVLTRSDKNRGKSDNVPSEEVVKKMKNYWRQLLNAKLITQRKFDNLTKAERGGLSELDKAGFIKRQLVETRQITKHVAQILDSRMNTKYDENDKLIREVKVITLKSKLVSDFRKDFQFYKVREINNYHHAHDAYLNAVVGTALIKKYPKLESEFVYGDYKVYDVRKMIAKSEQEIGKATAKYFFYSNIMNFFKTEITLANGEIRKRPLIETNGETGEIVWDKGRDFATVRKVLSMPQVNIVKKTEVQTGGFSKESILPKRNSDKLIARKKDWDPKKYGGFDSPTVAYSVLVVAKVEKGKSKKLKSVKELLGITIMERSSFEKNPIDFLEAKGYKEVKKDLIIKLPKYSLFELENGRKRMLASAGELQKGNELALPSKYVNFLYLASHYEKLKGSPEDNEQKQLFVEQHKHYLDEIIEQISEFSKRVILADANLDKVLSAYNKHRDKPIREQAENIIHLFTLTNLGAPAAFKYFDTTIDRKRYTSTKEVLDATLIHQSITGLYETRIDLSQLGGDSAGGGGSGLATTLERIEKNFVITDPRLPDNPIIFASDSFLQLTEYSREEILGRNCRFLQGPETDRA |

| | | |
|---|---|---|
| | | TVRKIRDAIDNQTEVTVQLINYTKSGKKFWNLFHLQPMRDQKGDVQYFIGVQLD GTEHVRDAAEREGVMLIKKTAENIDEAAKEL |
| 55 | dCas9-AsLOVx2 | MSPKKKRKVEASDKKYSIGLAIGTNSVGWAVITDEYKVPSKKFKVLGNTDRHSIK KNLIGALLFDSGETAEATRLKRTARRRYTRRKNRICYLQEIFSNEMAKVDDSFFH RLEESFLVEEDKKHERHPIFGNIVDEVAYHEKYPTIYHLRKKLVDSTDKADLRLIYL ALAHMIKFRGHFLIEGDLNPDNSDVDKLFIQLVQTYNQLFEENPINASGVDAKAIL SARLSKSRRLENLIAQLPGEKKNGLFGNLIALSLGLTPNFKSNFDLAEDAKLQLSK DTYDDDLDNLLAQIGDQYADLFLAAKNLSDAILLSDILRVNTEITKAPLSASMIKRY DEHHQDLTLLKALVRQQLPEKYKEIFFDQSKNGYAGYIDGGASQEEFYKFIKPILE KMDGTEELLVKLNREDLLRKQRTFDNGSIPHQIHLGELHAILRRQEDFYPFLKDN REKIEKILTFRIPYYVGPLARGNSRFAWMTRKSEETITPWNFEEVVDKGASAQSFI ERMTNFDKNLPNEKVLPKHSLLYEYFTVYNELTKVKYVTEGMRKPAFLSGEQKK AIVDLLFKTNRKVTVKQLKEDYFKKIECFDSVEISGVEDRFNASLGTYHDLLKIIKD KDFLDNEENEDILEDIVLTLTLFEDREMIEERLKTYAHLFDDKVMKQLKRRRYTG WGRLSRKLINGIRDKQSGKTILDFLKSDGFANRNFMQLIHDDSLTFKEDIQKAQV SGQGDSLHEHIANLAGSPAIKKGILQTVKVVDELVKVMGRHKPENIVIEMARENQ TTQKGQKNSRERMKRIEEGIKELGSQILKEHPVENTQLQNEKLYLYYLQNGRDM YVDQELDINRLSDYDVDAIVPQSFLKDDSIDNKVLTRSDKNRGKSDNVPSEEVVK KMKNYWRQLLNAKLITQRKFDNLTKAERGGLSELDKAGFIKRQLVETRQITKHVA QILDSRMNTKYDENDKLIREVKVITLKSKLVSDFRKDFQFYKVREINNYHHAHDA YLNAVVGTALIKKYPKLESEFVYGDYKVYDVRKMIAKSEQEIGKATAKYFFYSNIM NFFKTEITLANGEIRKRPLIETNGETGEIVWDKGRDFATVRKVLSMPQVNIVKKTE VQTGGFSKESILPKRNSDKLIARKKDWDPKKYGGFDSPTVAYSVLVVAKVEKGK SKKLKSVKELLGITIMERSSFEKNPIDFLEAKGYKEVKKDLIIKLPKYSLFELENGR KRMLASAGELQKGNELALPSKYVNFLYLASHYEKLKGSPEDNEQKQLFVEQHK HYLDEIIEQISEFSKRVILADANLDKVLSAYNKHRDKPIREQAENIIHLFTLTNLGAP AAFKYFDTTIDRKRYTSTKEVLDATLIHQSITGLYETRIDLSQLGGDSAGGGGSGL ATTLERIEKNFVITDPRLPDNPIIFASDSFLQLTEYSREEILGRNCRFLQGPETDRA TVRKIRDAIDNQTEVTVQLINYTKSGKKFWNLFHLQPMRDQKGDVQYFIGVQLD GTEHVRDAAEREGVMLIKKTAENIDEAAKELGSGSGSGSGLATTLERIEKNFVIT DPRLPDNPIIFASDSFLQLTEYSREEILGRNCRFLQGPETDRATVRKIRDAIDNQT EVTVQLINYTKSGKKFWNLFHLQPMRDQKGDVQYFIGVQLDGTEHVRDAAERE GVMLIKKTAENIDEAAKEL |
| 56 | dCas9-AsLOV2x3 | MSPKKKRKVEASDKKYSIGLAIGTNSVGWAVITDEYKVPSKKFKVLGNTDRHSIK KNLIGALLFDSGETAEATRLKRTARRRYTRRKNRICYLQEIFSNEMAKVDDSFFH RLEESFLVEEDKKHERHPIFGNIVDEVAYHEKYPTIYHLRKKLVDSTDKADLRLIYL ALAHMIKFRGHFLIEGDLNPDNSDVDKLFIQLVQTYNQLFEENPINASGVDAKAIL SARLSKSRRLENLIAQLPGEKKNGLFGNLIALSLGLTPNFKSNFDLAEDAKLQLSK DTYDDDLDNLLAQIGDQYADLFLAAKNLSDAILLSDILRVNTEITKAPLSASMIKRY DEHHQDLTLLKALVRQQLPEKYKEIFFDQSKNGYAGYIDGGASQEEFYKFIKPILE KMDGTEELLVKLNREDLLRKQRTFDNGSIPHQIHLGELHAILRRQEDFYPFLKDN REKIEKILTFRIPYYVGPLARGNSRFAWMTRKSEETITPWNFEEVVDKGASAQSFI ERMTNFDKNLPNEKVLPKHSLLYEYFTVYNELTKVKYVTEGMRKPAFLSGEQKK AIVDLLFKTNRKVTVKQLKEDYFKKIECFDSVEISGVEDRFNASLGTYHDLLKIIKD KDFLDNEENEDILEDIVLTLTLFEDREMIEERLKTYAHLFDDKVMKQLKRRRYTG WGRLSRKLINGIRDKQSGKTILDFLKSDGFANRNFMQLIHDDSLTFKEDIQKAQV SGQGDSLHEHIANLAGSPAIKKGILQTVKVVDELVKVMGRHKPENIVIEMARENQ TTQKGQKNSRERMKRIEEGIKELGSQILKEHPVENTQLQNEKLYLYYLQNGRDM YVDQELDINRLSDYDVDAIVPQSFLKDDSIDNKVLTRSDKNRGKSDNVPSEEVVK KMKNYWRQLLNAKLITQRKFDNLTKAERGGLSELDKAGFIKRQLVETRQITKHVA QILDSRMNTKYDENDKLIREVKVITLKSKLVSDFRKDFQFYKVREINNYHHAHDA YLNAVVGTALIKKYPKLESEFVYGDYKVYDVRKMIAKSEQEIGKATAKYFFYSNIM NFFKTEITLANGEIRKRPLIETNGETGEIVWDKGRDFATVRKVLSMPQVNIVKKTE VQTGGFSKESILPKRNSDKLIARKKDWDPKKYGGFDSPTVAYSVLVVAKVEKGK SKKLKSVKELLGITIMERSSFEKNPIDFLEAKGYKEVKKDLIIKLPKYSLFELENGR KRMLASAGELQKGNELALPSKYVNFLYLASHYEKLKGSPEDNEQKQLFVEQHK HYLDEIIEQISEFSKRVILADANLDKVLSAYNKHRDKPIREQAENIIHLFTLTNLGAP AAFKYFDTTIDRKRYTSTKEVLDATLIHQSITGLYETRIDLSQLGGDGSGSGSGSG LATTLERIEKNFVITDPRLPDNPIIFASDSFLQLTEYSREEILGRNCRFLQGPETDR ATVRKIRDAIDNQTEVTVQLINYTKSGKKFWNLFHLQPMRDQKGDVQYFIGVQL DGTEHVRDAAEREGVMLIKKTAENIDEAAKELSAGGGGSGLATTLERIEKNFVIT DPRLPDNPIIFASDSFLQLTEYSREEILGRNCRFLQGPETDRATVRKIRDAIDNQT EVTVQLINYTKSGKKFWNLFHLQPMRDQKGDVQYFIGVQLDGTEHVRDAAERE GVMLIKKTAENIDEAAKELGSGSGSGSGLATTLERIEKNFVITDPRLPDNPIIFASD SFLQLTEYSREEILGRNCRFLQGPETDRATVRKIRDAIDNQTEVTVQLINYTKSGK KFWNLFHLQPMRDQKGDVQYFIGVQLDGTEHVRDAAEREGVMLIKKTAENIDEA AKEL |

| | | |
|---|---|---|
| 57 | dCas9-AsLOV2x3-NLSx3 | MSPKKKRKVGSGPAAKRVKLDGSGPKKKRKVEASDKKYSIGLAIGTNSVGWAVI<br>TDEYKVPSKKFKVLGNTDRHSIKKNLIGALLFDSGETAEATRLKRTARRRYTRRK<br>NRICYLQEIFSNEMAKVDDSFFHRLEESFLVEEDKKHERHPIFGNIVDEVAYHEK<br>YPTIYHLRKKLVDSTDKADLRLIYLALAHMIKFRGHFLIEGDLNPDNSDVDKLFIQL<br>VQTYNQLFEENPINASGVDAKAILSARLSKSRRLENLIAQLPGEKKNGLFGNLIAL<br>SLGLTPNFKSNFDLAEDAKLQLSKDTYDDDLDNLLAQIGDQYADLFLAAKNLSDA<br>ILLSDILRVNTEITKAPLSASMIKRYDEHHQDLTLLKALVRQQLPEKYKEIFFDQSK<br>NGYAGYIDGGASQEEFYKFIKPILEKMDGTEELLVKLNREDLLRKQRTFDNGSIP<br>HQIHLGELHAILRRQEDFYPFLKDNREKIEKILTFRIPYYVGPLARGNSRFAWMTR<br>KSEETITPWNFEEVVDKGASAQSFIERMTNFDKNLPNEKVLPKHSLLYEYFTVYN<br>ELTKVKYVTEGMRKPAFLSGEQKKAIVDLLFKTNRKVTVKQLKEDYFKKIECFDS<br>VEISGVEDRFNASLGTYHDLLKIIKDKDFLDNEENEDILEDIVLTLTLFEDREMIEER<br>LKTYAHLFDDKVMKQLKRRRYTGWGRLSRKLINGIRDKQSGKTILDFLKSDGFA<br>NRNFMQLIHDDSLTFKEDIQKAQVSGQGDSLHEHIANLAGSPAIKKGILQTVKVV<br>DELVKVMGRHKPENIVIEMARENQTTQKGQKNSRERMKRIEEGIKELGSQILKEH<br>PVENTQLQNEKLYLYYLQNGRDMYVDQELDINRLSDYDVDAIVPQSFLKDDSIDN<br>KVLTRSDKNRGKSDNVPSEEVVKKMKNYWRQLLNAKLITQRKFDNLTKAERGG<br>LSELDKAGFIKRQLVETRQITKHVAQILDSRMNTKYDENDKLIREVKVITLKSKLVS<br>DFRKDFQFYKVREINNYHHAHDAYLNAVVGTALIKKYPKLESEFVYGDYKVYDV<br>RKMIAKSEQEIGKATAKYFFYSNIMNFFKTEITLANGEIRKRPLIETNGETGEIVWD<br>KGRDFATVRKVLSMPQVNIVKKTEVQTGGFSKESILPKRNSDKLIARKKDWDPK<br>KYGGFDSPTVAYSVLVVAKVEKGKSKKLKSVKELLGITIMERSSFEKNPIDFLEAK<br>GYKEVKKDLIIKLPKYSLFELENGRKRMLASAGELQKGNELALPSKYVNFLYLAS<br>HYEKLKGSPEDNEQKQLFVEQHKHYLDEIIEQISEFSKRVILADANLDKVLSAYNK<br>HRDKPIREQAENIIHLFTLTNLGAPAAFKYFDTTIDRKRYTSTKEVLDATLIHQSITG<br>LYETRIDLSQLGGDGSGSGSGSGLATTLERIEKNFVITDPRLPDNPIIFASDSFLQL<br>TEYSREEILGRNCRFLQGPETDRATVRKIRDAIDNQTEVTVQLINYTKSGKKFWN<br>LFHLQPMRDQKGDVQYFIGVQLDGTEHVRDAAEREGVMLIKKTAENIDEAAKEL<br>SAGGGGSGLATTLERIEKNFVITDPRLPDNPIIFASDSFLQLTEYSREEILGRNCR<br>FLQGPETDRATVRKIRDAIDNQTEVTVQLINYTKSGKKFWNLFHLQPMRDQKGD<br>VQYFIGVQLDGTEHVRDAAEREGVMLIKKTAENIDEAAKELGSGSGSGSGLATTL<br>ERIEKNFVITDPRLPDNPIIFASDSFLQLTEYSREEILGRNCRFLQGPETDRATVRK<br>IRDAIDNQTEVTVQLINYTKSGKKFWNLFHLQPMRDQKGDVQYFIGVQLDGTEH<br>VRDAAEREGVMLIKKTAENIDEAAKEL |
| 60 | Zdk2-VP64 | MGSGPKKKRKVAAAGGSGSGSMVDNKFNKEMLSARVEIYGLPNLNWGQRFAFI<br>SSLTDDPSQSANLLAEAKKLNDAQAPKGGGGSGGGGSAAAGSGRADALDDFD<br>LDMLGSDALDDFDLDMLGSDALDDFDLDMLGSDALDDFDLDMLGSGSGSSPKK<br>KRKVEAS |
| 61 | Zdk1-VP64 | MGSGPKKKRKVAAAGGSGSGSMVDNKFNKEKTRAGAEIHSLPNLNVEQKFAFI<br>VSLFDDPSQSANLLAEAKKLNDAQAPKGGGGSGGGGSAAAGSGRADALDDFD<br>LDMLGSDALDDFDLDMLGSDALDDFDLDMLGSDALDDFDLDMLGSGSGSSPKK<br>KRKVEAS |
| 62 | Zdk3-VP64 | MGSGPKKKRKVAAAGGSGSGSMVDNKFNKEVLVARQEIYWLPNLNWEQKFAFI<br>SSLTNDPSQSANLLAEAKKLNGAQPPKGGGGSGGGGSAAAGSGRADALDDFD<br>LDMLGSDALDDFDLDMLGSDALDDFDLDMLGSDALDDFDLDMLGSGSGSSPKK<br>KRKVEAS |
| 63 | Zdk3-p65 | MGSGPKKKRKVAAAGGSGSGSMVDNKFNKEVLVARQEIYWLPNLNWEQKFAFI<br>SSLTNDPSQSANLLAEAKKLNGAQPPKGGGGSGGGGSAAAGSGRAPTQAGEG<br>TLSEALLQLQFDDEDLGALLGNSTDPAVFTDLASVDNSEFQQLLNQGIPVAPHTT<br>EPMLMEYPEAITRLVTGAQRPPDPAPAPLGAPGLPNGLLSGDEDFSSIADMDFS<br>ALLSQISSGSGSGSSPKKKRKVEAS |
| 64 | Zdk1-VPR | MGSGPKKKRKVAAAGGSGSGSMVDNKFNKEKTRAGAEIHSLPNLNVEQKFAFI<br>VSLFDDPSQSANLLAEAKKLNDAQAPKGGGGSGGGGSAAAGSGRADALDDFD<br>LDMLGSDALDDFDLDMLGSDALDDFDLDMLGSDALDDFDLDMLINTSGGSGGG<br>SGGSSQYLPDTDDRHRIEEKRKRTYETFKSIMKKSPFSGPTDPRPPPRRIAVPS<br>RSSASVPKPAPQPYPFTSSLSTINYDEFPTMVFPSGQISQASALAPAPPQVLPQA<br>PAPAPAPAMVSALAQAPAPVPVLAPGPPQAVAPPAPKPTQAGEGTLSEALLQLQ<br>FDDEDLGALLGNSTDPAVFTDLASVDNSEFQQLLNQGIPVAPHTTEPMLMEYPE<br>AITRLVTGAQRPPDPAPAPLGAPGLPNGLLSGDEDFSSIADMDFSALLGSGSGS<br>RDSREGMFLPKPEAGSAISDVFEGREVCQPKRIRPFHPPGSPWANRPLPASLAP<br>TPTGPVHEPVGSLTPAPVPQPLDPAPAVTPEASHLLEDPDEDTSQAVKALREMA<br>DTVIPQKEEAAICGQMDLSHPPPRGHLDELTTTLESMTEDLNLDSPLTPELNEILD<br>TFLNDECLLHAMHISTGLSIFDTSLFGSGSGSSPKKKRKVEAS |
| 65 | Zdk3-VPR | MGSGPKKKRKVAAAGGSGSGSMVDNKFNKEVLVARQEIYWLPNLNWEQKFAFI<br>SSLTNDPSQSANLLAEAKKLNGAQPPKGGGGSGGGGSAAAGSGRADALDDFD<br>LDMLGSDALDDFDLDMLGSDALDDFDLDMLGSDALDDFDLDMLINTSGGSGGG |

| | | |
|---|---|---|
| | | SGGSSQYLPDTDDRHRIEEKRKRTYETFKSIMKKSPFSGPTDPRPPPRRIAVPS RSSASVPKPAPQPYPFTSSLSTINYDEFPTMVFPSGQISQASALAPAPPQVLPQA PAPAPAPAMVSALAQAPAPVPVLAPGPPQAVAPPAPKPTQAGEGTLSEALLQLQ FDDEDLGALLGNSTDPAVFTDLASVDNSEFQQLLNQGIPVAPHTTEPMLMEYPE AITRLVTGAQRPPDPAPAPLGAPGLPNGLLSGDEDFSSIADMDFSALLGSGSGS RDSREGMFLPKPEAGSAISDVFEGREVCQPKRIRPFHPPGSPWANRPLPASLAP TPTGPVHEPVGSLTPAPVPQPLDPAPAVTPEASHLLEDPDEETSQAVKALREMA DTVIPQKEEAAICGQMDLSHPPPRGHLDELTTTLESMTEDLNLDSPLTPELNEILD TFLNDECLLHAMHISTGLSIFDTSLFGSGSGSSPKKKRKVEAS |

**Supplementary Table 3: Targeted genomic loci.** A-F: different positions in the Promoter region of the targeted gene.

| Organism | Gene | Target | gRNA sequence | Genomic location | Reference |
|---|---|---|---|---|---|
| ***H. sapiens*** | **IL1RN** | Promoter A | TGTACTCTCTGAGGTGCTC | chr2:113875442-113875460 | [3] |
| | | Promoter B | ACGCAGATAAGAACCAGTT | chr2:113875291-113875309 | [3] |
| | | Promoter C | CATCAAGTCAGCCATCAGC | chr2:113875358-113875376 | [3] |
| | | Promoter D | GAGTCACCCTCCTGGAAAC | chr2:113875326-113875344 | [3] |
| ***H. sapiens*** | **MYOD** | Promoter A | CCTGGGCTCCGGGGCGTTT | chr11:17741056-17741074 | [3] |
| | | Promoter B | GGCCCCTGCGGCCACCCCG | chr11:17740969-17740987 | [3] |
| | | Promoter C | CTCCCTCCCTGCCCGGTAG | chr11:17740897-17740915 | [3] |
| | | Promoter D | AGGTTTGGAAAGGGCGTGC | chr11:17740837-17740855 | [3] |
| ***H. sapiens*** | **OCT4** | Promoter A | ACTCCACTGCACTCCAGTCT | chr6:31138711-31138730 | [4] |
| | | Promoter B | TCTGTGGGGGACCTGCACTG | chr6:31138643-31138662 | [4] |
| | | Promoter C | GGGGCGCCAGTTGTGTCTCC | chr6:31138613-31138632 | [4] |
| | | Promoter D | ACACCATTGCCACCACCATT | chr6:31138574-31138593 | [4] |
| ***M. musculus*** | **Kir2.1** | Promoter A | GCAGCCACGTGCGCTCCGAT | chr11:110956901-110956920 | This study |
| | | Promoter B | TCCTCCTGATGGCGGTTCCC | chr11:110956867-110956886 | This study |
| | | Promoter C | ATCTCTACACCCGCGGACCC | chr11:110956836-110956855 | This study |
| | | Promoter D | AGCACCGGCAAGCAGTGTCT | chr11:110957028-110957047 | This study |
| | | Promoter E | CATGATCCTGTACCAGCAAC | chr11:110957139-110957158 | This study |
| | | Promoter F | GGAAGGCTGCGCGCTGGCCA | chr11:110957007-110957026 | This study |
| ***M. musculus*** | **Neurog2** | Promoter A | CGGGCAGATCTGATTGTTTTC | chr3:127426729-127426748 | This study |
| ***M. musculus*** | **Arc** | Promoter A | ATTGGCGGCGCCCAGGCTCC | chr15:74544493-74544512 | This study |
| | | Promoter B | GCCTTCCGCGCCTACTCGCT | chr15:74544553-74544572 | This study |
| | | Promoter C | CCCGGTGGGAGGCGCGCAGC | chr15:74544635-74544654 | This study |
| | | Promoter D | GTTCGGAGCCGCAGCACCGA | chr15:74544229-74544248 | This study |
| | | Promoter E | CACTCGCTAAGCTCCTCCGG | chr15:74544327-74544346 | This study |

**Supplementary Table 4: RT-qPCR primers used in this study.**

| Gene | Forward primer | Reverse primer | Reference |
|---|---|---|---|
| **GAPDH** | CAATGACCCCTTCATTGACC | TTGATTTTGGAGGGATCTCG | [3] |
| **IL1RN** | GGAATCCATGGAGGGAAGAT | TGTTCTCGCTCAGGTCAGTG | [3] |
| **MYOD** | GACGGCATGATGGACTACAG | GATGCTGGACAGGCAGTCT | This study |
| **Oct4** | CGAAAGAGAAAGCGAACCAGTATCGAGAAC | CGTTGTGCATAGTCGCTGCTTGATCGC | [4] |
| **Kir2.1** | ACTTGCTTCGGCTCATTCTCT | CCAGAGAACTTGTCCTGTTGCT | This study |
| **Arc** | GAACCTCAACTTCCGGGGAT | ACTGGTATGAATCACTGGGGG | This study |
| **Neurog2** | AACTCCACGTCCCCATACAG | GAGGCGCATAACGATGCTTCT | This study |
| **Actin** | TGTTACCAACTGGGACGACA | ACCTGGGTCATCTTTTCACG | This study |
